# Genetic variation underlying plasticity in physiological traits mediates response to climate in a *Populus* hybrid zone

**DOI:** 10.64898/2026.05.13.724884

**Authors:** Michelle Zavala-Paez, Alayna Mead, Baxter Worthing, Sara K Klopf, Stephen Keller, Jason Holliday, Matthew C. Fitzpatrick, Jill A Hamilton

## Abstract

Phenotypic plasticity can buffer the potential fitness consequences of environmental change, yet limited understanding of its genetic basis constrains its application to predicting population response to future climates. Hybrid zones provide powerful systems to study the genetic basis of plasticity because admixture can create novel allele combinations that generate new reaction norms for selection to act upon. Here, we combine two clonally replicated common gardens of *Populus trichocarpa* × *P. balsamifera* genotypes with whole-genome resequencing to identify genetic variation underlying plasticity for physiological traits. Admixed genotypes exhibited broader, and in some cases novel, reaction norms relative to parental genotypes, particularly for key stomatal traits. Admixture mapping of genotype-specific reaction norms identified ten candidate genes on chromosome 15, including *TWIST*, associated with plasticity in adaxial stomatal occurrence and density. Using random forest models, we projected allele-specific responses to climate warming within the hybrid zone to link genetic variation in plasticity with predicted warming. Random forest models forecast that future climates would favor *P. trichocarpa* alleles at *TWIST*, while *P. balsamifera* alleles would be maintained in heterozygous genotypes. These results suggest that hybridization can expand reaction norms and maintain genetic variation that may facilitate rapid phenotypic response needed to adapt to climate change.

## Introduction

In a changing climate, plant populations must either migrate to favorable habitats, adapt, or persist locally through mechanisms that buffer the fitness costs associated with rapid environmental change (Aitken et al. 2008; Nicotra et al. 2010; Isabel et al. 2020). Phenotypic plasticity, the capacity of a genotype to modify phenotypic expression in response to varying environmental conditions, can provide the buffer needed to mediate rapidly changing conditions (Bradshaw et al. 1965; Scheiner 1993; Arnold et al. 2019; Schneider 2022). Importantly, plasticity can be heritable, and recent work has begun to uncover its genetic basis for key agronomic traits (e.g. flowering time and biomass) improving predictions for phenotypic expression across environments (Jin et al. 2023; Prohaska et al. 2024a; Alseekh et al. 2025). Plasticity is particularly important for physiological traits, which often adjust phenotypic expression in response to the environment, impacting plant fitness (Sultan 2000; Solé-Medina et al. 2022; Ramírez-Valiente et al. 2025). Plasticity for traits related to photochemistry, stomatal regulation, water-use efficiency, and nutrient uptake can mediate how effectively plants acquire and allocate nutrients, adjusting their phenotypic optima in response to environmental change (Hetherington and Woodward 2003; Poorter et al. 2009; Buckley 2019; Navarro et al. 2022). Importantly, the magnitude and direction of plasticity can differ among genotypes, indicating genetic variation for trait plasticity where natural selection can act upon (Scheiner 1993; Matesanz and Ramírez-Valiente 2019; Schneider 2022). However, the evolution of trait plasticity depends on both its heritability and the strength of selection (Napier et al. 2023). Therefore, identifying the genetic variation underlying plasticity and quantifying its evolutionary potential is essential for predicting the capacity of plant populations to respond to and track environmental change.

Reaction norms, which capture the shape and magnitude of phenotypic expression across an environmental gradient for a single genotype, provide a classic framework to describe the ability of a plant to modify its phenotype in response to environmental change (Scheiner 1993; Arnold et al. 2019). Genotypic variation in the reaction norm can arise from the evolution of genetic variation in how environmental cues are perceived and translated into different phenotypes (Jin aet al., 2023). Interspecific gene flow between species with contrasting environmental niches, which may have evolved genetic differences, can recombine regulatory and coding alleles that functionally underlie plasticity modifying the magnitude and direction of reaction norms (Hamilton and Miller 2016; Liu et al. 2021; Jin et al. 2023; Schwartz et al. 2024; Runemark et al. 2025). As a result, admixed genotypes can exhibit reaction norms outside or intermediate to the range of parental species or resemble one or other parents (Rieseberg et al. 1999; Liu et al. 2021; Schwartz et al. 2024). Reaction norms that result in transgressive phenotypic expression may allow admixed genotypes to persist in, or move into, environmental conditions not previously experienced by either parent, enhancing persistence under rapid environmental change (Schwartz et al. 2024). Therefore, understanding how genotypes with different genomic ancestries modify phenotypic expression across environments is critical for evaluating how hybridization influences evolutionary potential under ongoing environmental change.

Despite advances in identifying the genetic variation underlying traits critical to adaptation, the mechanisms that allow a single genotype to express different phenotypes across varying environments remain poorly understood. Empirical studies of the genetic basis of plasticity have largely focused on crops and their wild relatives (Liu et al. 2021; Fu and Wang 2023; Jin et al. 2023; Prohaska et al. 2024b; Alseekh et al. 2025). Within these systems, key regulatory elements involved in temperature signaling and environmental responsiveness have been identified (e.g. *ZmTPS14*, *PIF4*, *RCC1*, and *SPL*; Jin et al., 2023; Alseekh et al., 2025). Notably, some loci are genetically correlated, associated with both trait variation and trait plasticity suggesting that the same genomic region can shape trait expression and the regulation of that expression across different environments (Li et al. 2018; Jin et al. 2023; Alseekh et al. 2025). Under this scenario, reaction norms may evolve indirectly as a byproduct of selection that acts directly on trait variation, or vice versa (Via 1993; Lafuente et al. 2024). However, in other cases, the genetic architecture underlying trait plasticity appears to be largely independent, suggesting distinct regulatory mechanisms for traits and their plasticity (Jin et al. 2023). In such cases, genetic variation underlying a trait and its expression could evolve independently (Lafuente et al. 2024). Identifying the genetic variation underlying trait variation and its plasticity will allow us to evaluate whether responses to selection are constrained by shared genetic basis or facilitated by the independent evolution of these components under climate change.

*Populus* (L.) provides an ideal system for studying the genetic variation underlying plasticity as clonal propagation ensures identical genotypes can be planted across varying environments, allowing separation of environmental response from genetic effects underlying trait expression (Taylor 2002; Tuskan et al. 2006; Hidalgo et al. 2010). For *Populus*, trait expression and plasticity have been associated with climatic gradients, highlighting the role of extrinsic selection in their evolution across environments (Mckown et al. 2014; Evans et al. 2016; Viger et al. 2016; Liu and El-Kassaby 2019; Thibault et al. 2020; Cooper et al. 2022; Eisenring et al. 2022). These studies suggest that there is substantial genetic variation underlying trait variation and plasticity in *Populus*. Within the genus, *P. trichocarpa* (Torr. & Gray) and *P. balsamifera* (L.) are sister taxa adapted to contrasting environments that hybridize naturally, providing an ideal system for examining how interspecific gene flow influences the evolution of traits and their plasticity (Levsen et al. 2012; Geraldes et al. 2014; Richardson et al. 2014; Bolte et al. 2024). *P. trichocarpa* typically persists across moderate maritime environments characterized by moist and humid conditions (Suarez-Gonzalez et al. 2018; Bolte et al. 2024; Mead et al. 2026a). In contrast, *P. balsamifera* is a widely distributed boreal species adapted to seasonal extremes, including warm, dry summers and moist cold winters (Richardson et al. 2014; Suarez-Gonzalez et al. 2018; Mead et al. 2026a). The two species hybridize through the Rocky Mountains, forming natural hybrid zones that span a climatic transition from maritime to continental climates where a broad range of recombinant genotypes have been observed (Levsen et al. 2012; Geraldes et al. 2014; Bolte et al. 2024). Previous work in this hybrid zone has revealed substantial genetic variation for plasticity, with hybrids exhibiting intermediate reaction norms relative to the parental species for leaf phenology and physiological traits when planted in warmer environments (Mead et al. 2026b). The broad range of hybrid genotypes and substantial genetic variation in trait plasticity make this hybrid zone an ideal natural system to study the genetic basis of plasticity. In addition, this hybrid zone provides an opportunity to predict how allele-specific variation underlying plasticity may respond to environmental change

Here, we use clonally replicated *Populus* genotypes, including *P. trichocarpa*, *P. balsamifera*, and their hybrids, planted across two contrasting environments in North America to identify the genetic variation underlying plasticity for physiological traits important for adaptation. Specifically, we ask 1) to what extent do physiological traits exhibit genetic variation in plasticity across environments, quantified by genotype-by-environment interactions (G×E)? 2) to what extent does hybridization influence trait variation and its plasticity? 3) to what extent does genetic variation underlying trait variation regulate trait plasticity? and 4) how is genetic variation underlying phenotypic plasticity distributed across the landscape today, and how might selection under future climatic conditions shape its distribution in the future? These analyses reveal how interspecific gene flow shapes phenotypic responses across environments for *Populus* and identify the genetic variation underlying trait variation and its plasticity, and its potential to evolve in response to changing environmental conditions.

## Material and Methods

### Sample collection and library preparation

During the winter of 2020, dormant vegetative cuttings were collected for each of 574 genotypes across six natural contact zones spanning a broad latitudinal gradient between *Populus trichocarpa × P. balsamifera* (Figure 1; details in Bolte et al., 2024). Cuttings were clonally propagated in a greenhouse at Virginia Tech (Blacksburg, VA, USA) under controlled conditions (24 °C day/15.5 °C night, no supplemental lighting). Following establishment, ∼100 mg of fresh leaf tissue was harvested from each genotype for DNA extraction and whole-genome resequencing. Methodology associated with propagation, DNA extraction, library preparation, sequencing, and quality control is described in detail in Bolte et al. (2024) and in the Supplementary Methods here. After filtering for quality control, Bolte et al. (2024) retained 29,663,130 biallelic SNPs. In this study, we further filtered loci with a minor allele frequency <5% loci with >10% missing data and removed 30 genotypes that were identified as other *Populus* species based on genetic clustering results from Bolte et al 2024, resulting in a final dataset of 7,167,726 biallelic SNPs and 544 genotypes for downstream analyses.

**Figure 1.**
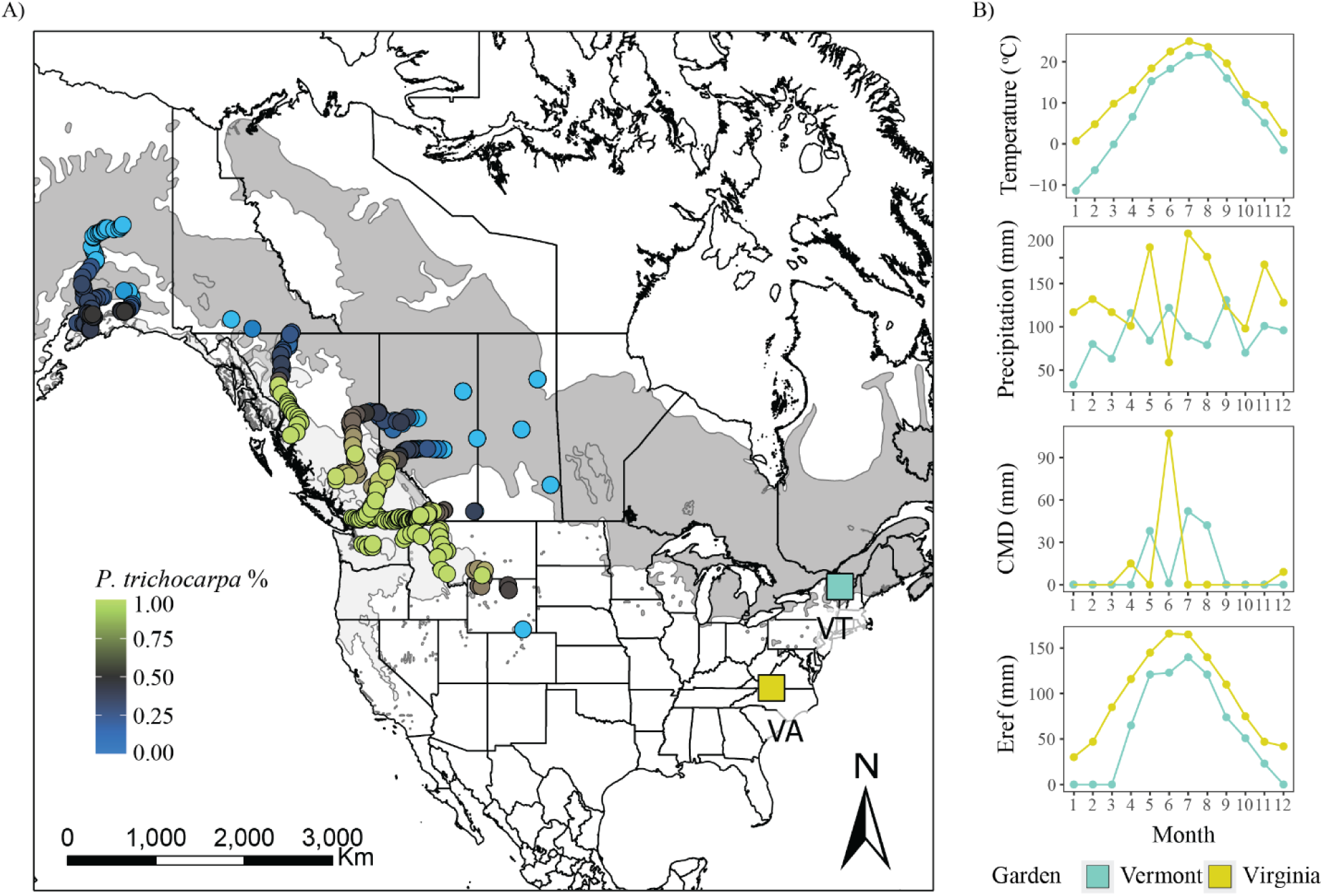
(A) Sampling locations across six *Populus trichocarpa × P. balsamifera* contact zones. Circles represent individual genotypes, colored by the proportion of *P. trichocarpa* nuclear ancestry based on whole-genome resequencing. Common garden experiments established in Vermont (VT, teal square) and Virginia (VA, yellow square) are shown. (B) For 2022, monthly means of temperature (°C), precipitation (mm), Hargreaves climatic moisture deficit (CMD; mm), and Hargreaves reference evaporation (Eref; mm) are provided for common garden locations.

### Common gardens design and phenotyping

In March 2020, the clonally propagated *Populus* cuttings were planted in two environments using a randomized complete block design, including three replicates per each of the 544 genotypes within each environment (Figure 1A). Common garden experiments were planted in Stuart, Virginia (36° 37’ N, 80° 09’ W; elevation 351 m), and Burlington, Vermont (44° 26’ N, 73° 11 W; elevation 130 m), which exhibit differences in mean monthly temperature (°C), precipitation (mm), Hargreaves climatic moisture deficit (mm), and Hargreaves reference evaporation (mm; Figure 1B). This design enabled quantification of genetic, environmental, and genotype-by-environment interactions underlying trait variability. In the summer of 2022, we quantified a suite of physiological traits across the two common garden environments (Details in Supplementary Methods; Table 1, Zavala-Paez et al. 2025; Zavala-Paez et al. 2026). These traits included: (i) leaf morphological traits: leaf dry mass (g), leaf area (m^2^), and leaf mass per area (gm^-2^), (ii) gas exchange traits: stomatal conductance (mol m^−2^ s^−1^) and intrinsic water-use efficiency (‰); (iii) photochemistry traits: quantum efficiency in light, electron transport rate (mol m^−2^ s^−1^), minimum fluorescence in light, and maximum fluorescence in light; (iv) stomatal patterning traits: adaxial (µm) and abaxial (µm) stomatal pore length, adaxial (µm) and abaxial (µm) stomatal density, stomatal ratio, and adaxial stomatal occurrence; and (iv) nutrient-use traits: leaf nitrogen content (%), leaf nitrogen isotopes (‰), leaf carbon content (%), and the leaf carbon-to-nitrogen ratio.

**Table 1.**
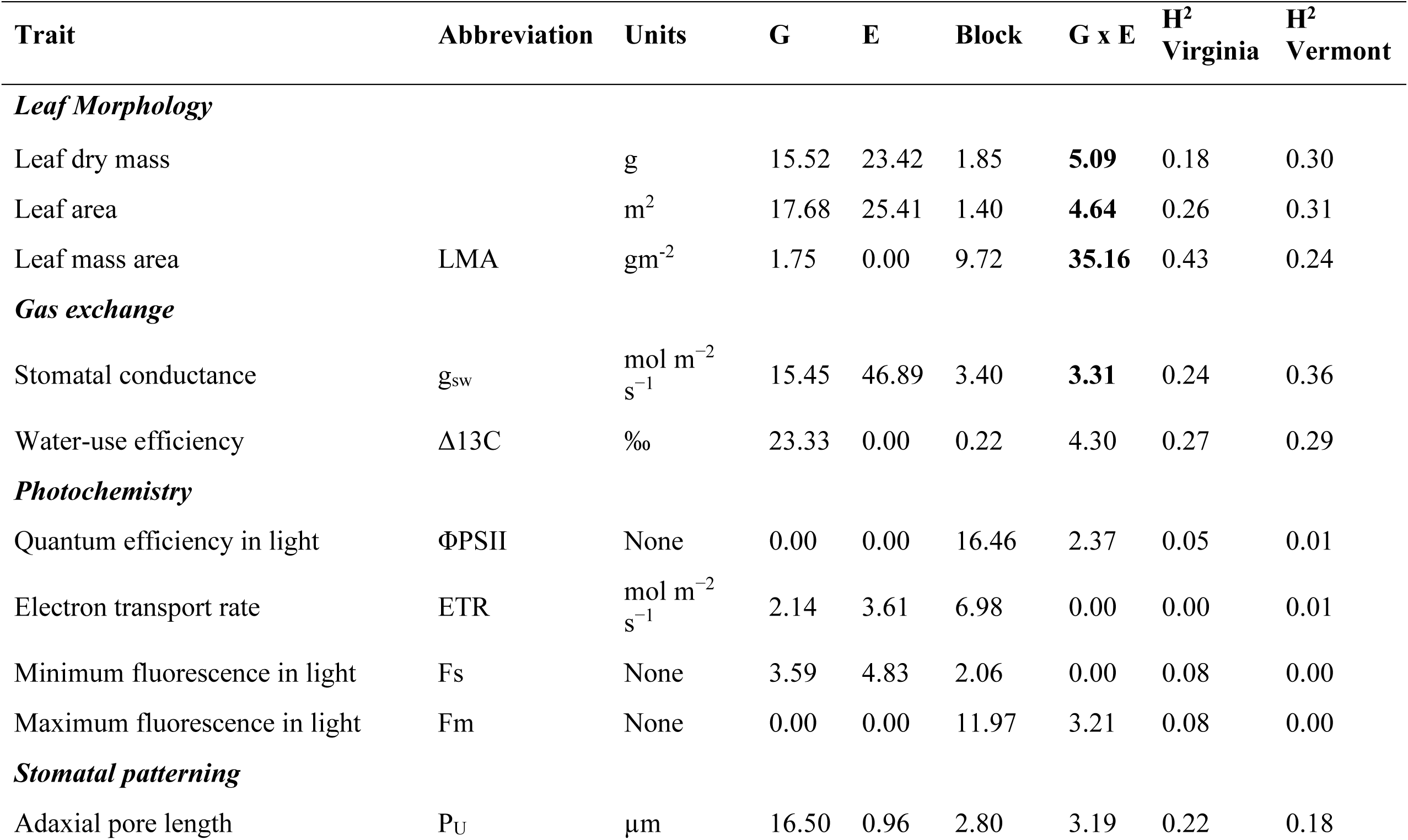
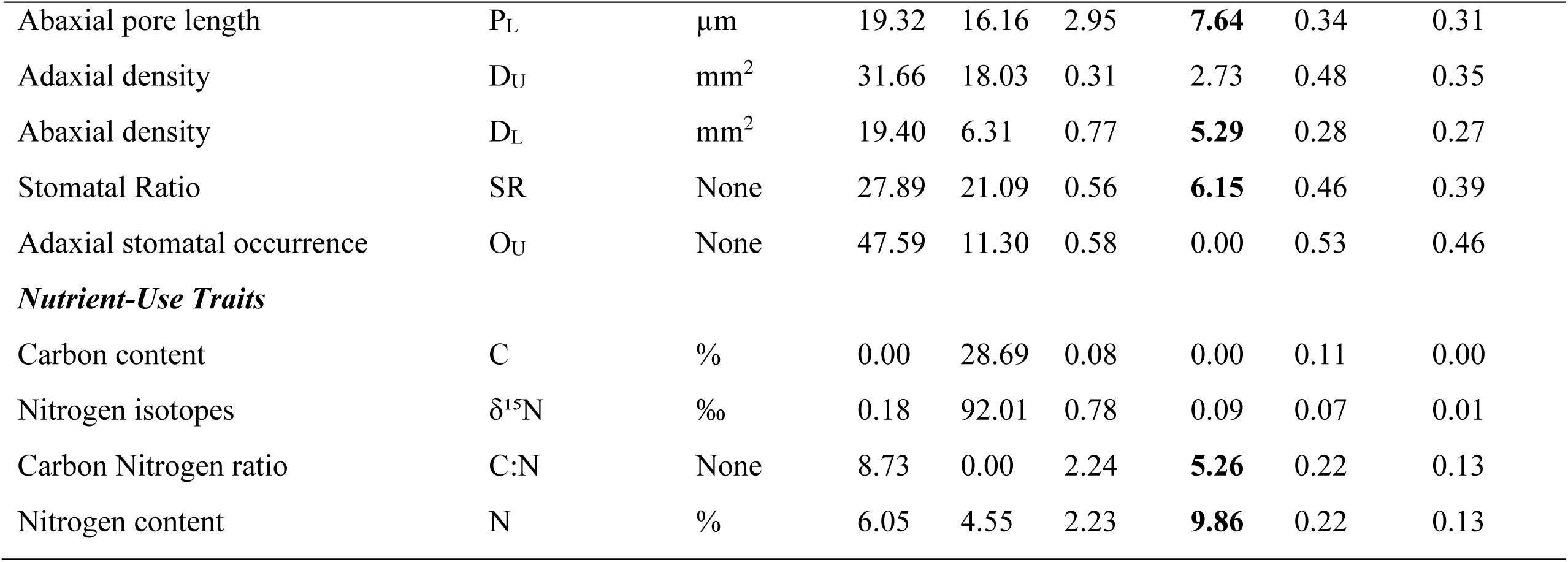
Physiological traits included in this study, grouped by functional class. Abbreviations and measurement units are provided for each trait. Broad-sense heritability (H^2^) is provided for each trait within each common garden environment (equation 2), along with the percent of phenotypic variance explained by genotype (G), environment (E), and their interaction (G×E, equation 1). Traits with statistically significant G×E interactions, based on likelihood ratio tests (LRT) comparing models with and without the interaction term, are in bold.

### Partitioning the genotype (G), environment (E) and G х E contributions to physiological trait variation

To estimated genetic variation for trait plasticity, quantified by genotype-by-environment interactions (G×E), we estimated the proportion of phenotypic variance explained by genotype (G), environment (E) and G х E for each trait using equation (1), with all models fitted using *lme*4 package in R version 4.3.1 (R Core Team, 2021):

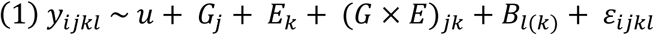

where *y*_*ijkl*_ represents the clonal individual *i* for the genotype *j* planted in the common garden *k*. The term *u* represents the overall model mean, *G*_*j*_ represents the random effect of genotype, *E*_*k*_ is the random effect of common garden *k* and (*G* × *E*)_*jk*_ is the random interaction effect between genotype and common garden, and *B*_*l*(*k*)_is the random effect of block *l* nested within common garden *k* and *ε*_*ijkl*_ is the residual variance. For adaxial stomata occurrence, which was measured as presence or absence, we used the *glmer* function with a binomial link function in R. After fitting the models, we extracted the variance components using the *VarCorr* function in R. To calculate the proportion of phenotypic variance explained for each component, we divided the estimate of variance for each term by the total phenotypic variance observed. Then, likelihood ratio tests were used to test whether the contribution of the G × E to phenotypic variation was statistically significant. Specifically, we fitted a reduced version of model (1) by removing the random effect of G × E, and compared it to the full model (1) using a likelihood ratio test implemented with the *anova* () function in R.

### Estimating broad sense heritability for physiological traits

To assess the proportion of variance in phenotypic traits due to genetic factors, we estimated broad-sense heritability (*H*^2^) for each trait separately within each common garden. Variance components, including genetic variance (*V_G_*) and total phenotypic variance (*V_P_*), were estimated by fitting a linear mixed-effects model for each garden using equation (2):

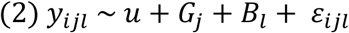

Where *y*_*ijl*_ is the observed trait value for the clonal individual *i* for the genotype *j* and block *l*. *G*_*j*_is the random effect of genotype representing the genetic effects (*V_G_*), *B*_*l*_ is the random effect of block representing environmental effects (*V_E_*) and *ε*_*ijl*_ is the residual error (*V_ɛ_*). Variance components from the linear mixed-effects model were extracted using *VarCorr* in R. *H^2^* was calculated using equation (3):

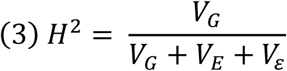

For adaxial stomatal occurrence, which is a binary variable, we fit a logistic mixed-effects model with the same random effects as in equation 2 using the *glmer* function in R (Bates et al., 2015).

### Estimating physiological trait variation and trait plasticity and their relationship with genomic ancestry

To quantify the association between ancestry, trait variation, and its plasticity, we first estimated genotype-level trait values using best linear unbiased predictors (BLUPs) and quantified trait plasticity for each genotype.

BLUPs, which represent the genetic value of each trait for each genotype (hereafter referred to as trait variation), were estimated using linear mixed models according to equation (4):

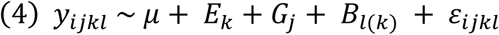

Where *y*_*ijkl*_ is the observed trait value for the clonal replicate *i*, *E*_*k*_ is the fixed effect of the common garden, *G*_*j*_ is the random effect of genotype *j*, *B*_*l*(*k*)_ is the random effect of block *l* nested with garden *k* and *ε*_*ijkl*_ is the residual term. The random effects of genotype were extracted from the model and used as estimates of genotype-level trait values. By accounting for environment and block as random effects in model (4), we obtained genotype-level estimates for each trait reflecting genetic differences among individuals.

To quantify trait plasticity, genotype reaction norms were estimated using linear mixed models with the function *lmer* from the lme4 package in R using equation (5):

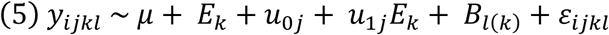

Where *y*_*ijkl*_ is the observed trait value for the clonal individual *i*, *E*_*k*_ is the fixed effect of the common garden site, *u*_0*j*_ is the random intercept for genotype *j*, *u*_1*j*_ is the random slope for the effect of garden on the genotype *j, B*_*l*(*k*)_ is the random effect of the block *l* nested with garden *k*. *ε*_*ijkl*_ represents the residual error of the model. The random slope for genotype *j*, which indicates the genotype-specific reaction norm for plasticity, was extracted and used as a metric of plasticity for subsequent analyses. A logistic mixed-effects model was used for stomatal occurrence to account for its binomial distribution.

To test whether genotypes with higher trait values also exhibit greater or reduced plasticity, we performed Spearman rank correlations between trait variation (BLUPs) and trait plasticity (genotype-specific reaction norms) in R. Significant correlations suggest a genetic correlation where trait variation and its plasticity are non-independent.

To test whether genomic ancestry predicts trait variation and trait plasticity, each response variable was modeled independently as a function of ancestry using regression models in R. Ancestry was defined as admixture proportions (*K*=2) using estimates from Mead et al (2026), with values ranging from 0 to 1 representing *P. trichocarpa* ancestry. We modeled the relationship between ancestry and trait variation or trait plasticity using equation (6):

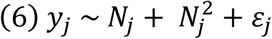

Where *y*_*j*_ represents the trait variation or plasticity for genotype *j*, *N*_*j*_ represents the ancestry of genotype *j* and 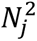 represent the quadratic term of ancestry to account for non-linear ancestry effects on the response variables. A reduced model including only the linear ancestry term *N*_*j*_ was also fit to evaluate whether the quadratic term improved model fit. Linear and quadratic models were compared using AIC, and the model with the lowest AIC was selected as the best-fit model for each trait.

### Admixture mapping to identify the genetic basis for physiological traits and their plasticity

To identify and compare the genetic variation underlying physiological traits and their plasticity, we conducted GWAS by admixture mapping to integrate local ancestry inference with genotype–phenotype associations. Prior to analysis, genomic data were phased and imputed for missingness using BEAGLE (beagle.06Aug24.a91.jar) for 544 genotypes (Browning et al. 2021). Local ancestry was estimated using Loter (Dias-Alves et al. 2018), which infers the ancestral origin of genomic regions in admixed individuals based on phased reference haplotypes. For local ancestry estimation, reference parental haplotypes were identified based on admixture proportions (*K* = 2, (Mead et al. 2026a). Genotypes with ancestry proportions (*Q*) less than 0.02 were designated as *P. balsamifera* (*n =* 90), those with *Q* greater than 0.98 as *P. trichocarpa* (*n*= 131, Zavala-Paez et al. 2026). These phased reference genotypes served to infer locus-specific ancestry across 323 admixed individuals using Loter. To associate locus-specific ancestry with trait variation and trait plasticity, we applied separate univariate linear mixed models for each trait and its plasticity using GEMMA v0.94.1 (Zhou and Stephens 2014). A kinship matrix and global ancestry (*K* =2) were included as covariates to account for relatedness and population structure. To correct for multiple testing, we used an admixture burden approach to estimate the effective number of independent tests, resulting in a significance threshold of log_10_(0.05/5,055) = 5.004. Full model specifications, admixture burden calculations, and diagnostic procedures are provided in the Supplementary Methods.

To account for the correlation between traits and their plasticity, we conducted an additional admixture mapping analysis in which plasticity for each trait was scaled according to genotype-specific trait means (i.e., scaled relative to the intercept). The full GWAS pipeline described above was then repeated for each trait using standardized plasticity values as the response variable.

### Forecasting shifts in genetic variation underlying plasticity

To model potential changes in local ancestry needed for plasticity in response to future climate projections, we used candidate genes associated with plasticity identified through admixture mapping. Twenty-five annual climate variables averaged between 1961–1991 sourced from ClimateNA (Wang et al. 2016) were used to represent historical climate conditions associated with genotype origin and to define the climate-of-origin parameter space used to project future shifts in local ancestry under climate change. We used the period between 1961-1991 as it likely reflects the climatic averages associated with selection influencing standing genetic variation for genotypes sourced within this study.

Future climate projections were modeled through ClimateNA (Mahony et al. 2022) using the Coupled Model Intercomparison Project Phase 6 (CMIP6). Future climatic conditions were modeled based on eight General Circulation Models (GCMs): ACCESS-ESM1-5, CNRM-ESM2-1, EC-Earth3, GFDL-ESM4, GISS-E2-1-G, MIROC6, MPI-ESM1-2-HR, and MRI- ESM2-0. These models were selected to capture variability among climate model structures while maintaining performance under historical conditions across North America (Mahony et al. 2022). Four Shared Socioeconomic Pathways (SSP1-2.6, SSP2-4.5, SSP3-7.0, and SSP5-8.5) were evaluated to capture a gradient of potential greenhouse gas trajectories, from strong mitigation (SSP126) to high-emission scenarios (SSP585). Climate projections were summarized for three 30-year intervals (2011–2040, 2041–2070, and 2071–2100), representing early-, mid-, and late-century periods. To compare projected future climates with historical climate conditions, paired *t*-tests were performed for each climate variable using each genotype’s climate of origin as replicates (Klein et al. 2025). *t*-statistics were used to indicate departures between future climates and historical conditions for a given SSP projection and time interval.

We used a random forest (RF) classification model, implemented in the *caret* package in R (Kuhn 2008), to predict potential shifts in genetic variation at plasticity-associated candidate genes under future climates. Random forest models were used because they are non-parametric, robust to multicollinearity among predictors, and well suited for capturing complex, non-linear relationships between genetic and climate variation. The model was trained using 25 annual historical climatic variables and local ancestry class assignment for candidate genes associated with plasticity derived from local ancestry analysis (homozygous *P. trichocarpa*, homozygous *P. balsamifera*, or heterozygous). To minimize bias across local ancestry classes, random down-sampling for the 544 genotypes was used to balance representation across homozygous *P. trichocarpa*, heterozygous, and homozygous *P. balsamifera* ancestry classes resulting in 115 genotypes per class. Each RF model consisted of 1,000 trees tested with 3, 5, 7, and 9 climate predictors randomly sampled at each split. Model performance was evaluated using 10-fold cross-validation, with overall accuracy and Cohen’s κ used to assess agreement between predicted and observed ancestry classifications beyond chance expectations. The trained RF model was applied to the future climate scenarios (all combinations of GCM × SSP × time interval) to project the probability of ancestry for candidate genes across climates. To account for stochastic variation, the RF model was bootstrapped 100 times, each initiated with a different random seed and a newly balanced training dataset generated by resampling with replacement. Probabilities predicted following resampling were averaged for each genotype and future climate scenario to estimate the local ancestry class for the candidate gene locus across individual genotypes under each genotype–climate scenario. The local ancestry class with the highest mean probability was assigned as the expected ancestry class for that locus under changing conditions. Finally, to estimate directional changes in the frequency of allelic variation associated with environmental response, we compared projected genotype-level local ancestry class assignments at candidate gene loci between historical and future climate scenarios.

## Results

### Physiological traits exhibit genetic variation in plasticity

Genetic variation for plasticity differed among traits, as reflected by variation in the proportion of phenotypic variance explained by genotype-by-environment interactions (G×E) (Figure S1). After testing for the significance of G×E, we found that G×E accounted between 3.31–35.16% of the variance observed for 9 out of 19 traits spanning leaf morphology, stomatal patterning, gas exchange, and nutrient-use trait classes (Table 1). Among traits, leaf mass area exhibited the greatest amount of genetic variation in reaction norms, with G×E explaining 35.16% of the variance observed. These results indicate that genetic variation underlying plasticity is present across multiple trait classes.

Differences across genotypes (G) were the primary contributor to trait variation, except those associated with photochemistry and nutrient use (Figure S1). The influence of genotype was particularly strong for the stomatal patterning trait class, as genotype explained 16.50–47.59% of the trait variance observed (Table 1). Environmental differences among common garden experiments (E) explained substantial variance in traits associated with nutrient-use, with environment accounting for 92.01% of the variation observed for nitrogen isotopes across environments. Strong block effects within garden were observed for photochemistry traits, suggesting that micro-environmental variation within gardens explained the majority of variance for these traits. For the majority of traits, the largest proportion of total phenotypic variance was explained by genotypic effects, whereas for select trait groups (e.g., leaf morphology), G×E became an important predictor of trait variation (Figure S1).

### Heritability of physiological traits across environments

On average, broad-sense heritability varied widely across different phenotypic traits, however, heritability values were largely consistent across environments, indicating limited environmental dependence of genetic effects (Table 1). Stomatal patterning traits exhibited the highest trait heritability across both gardens (*H^2^* Vermont = 0.18–0.46; *H^2^* Virginia = 0.22–0.53), followed by leaf morphological traits (*H^2^* Vermont = 0.24–0.31; *H^2^*Virginia = 0.18–0.43) indicating that variation in these traits has a substantial genetic component that is maintained irrespective of the environment. Gas exchange traits exhibited moderate heritability (*H^2^*Vermont = 0.29–0.36; *H^2^* Virginia = 0.24–0.27), whereas nutrient-use traits were generally lower (*H^2^* Vermont = 0.00- 0.13; *H^2^* Virginia = 0.07–0.22). Light-use efficiency traits had *H^2^* values near zero for all measured traits indicating that variation in photosynthetic efficiency is likely driven mainly by environmental factors.

### Hybridization influences the magnitude and direction of reaction norms across trait classes

Trait variation and trait plasticity were strongly correlated (Figure S2), with genomic ancestry influencing trait variation but also the magnitude and direction of reaction norms across trait classes (Figure 2). Overall, genotypes with higher *P. trichocarpa* ancestry exhibited greater leaf dry mass (*slope* = 0.45 g, *p* < 0.01, Figure S3), larger leaf area (*slope* = 0.56 m^2^, *p* < 0.01), higher intrinsic water-use efficiency (*slope* = 1.07 ‰, *p* < 0.05), greater adaxial stomatal density (*slope* = -2.59 mm^2^, *p* < 0.05), higher stomatal ratio (*slope* = -0.21 *p* < 0.001), and more adaxial stomatal occurrence (*slope* = 5.64, *p* < 0.001). Genotypes with higher *P. balsamifera* ancestry had higher leaf mass per area (*slope* = -4.32 gm^-2^, *p* > 0.001, Figure 2A), stomatal conductance (*slope* = -0.73 mol m^−2^ s^−1^, *p* > 0.001), and photochemical efficiency (Table S1). Genotypes at intermediate ancestries (hybrids) generally exhibited intermediate values for leaf morphological and photochemical traits. However, some traits had nonlinear relationships with ancestry (Table S1), with traits changing gradually at intermediate ancestry and more sharply toward parentals. For stomatal traits, hybrids had larger pore lengths than either parental species across both leaf surfaces, whereas abaxial stomatal density was reduced in hybrids relative to parental genotypes. Hybrids had higher nitrogen content and nitrogen isotope values and lower carbon-to-nitrogen ratios, indicating potential differences in nitrogen acquisition or allocation relative to parental types. Genotypes with intermediate ancestry had the highest stomatal conductance, but it declined with increasing *P. trichocarpa* ancestry, while water-use efficiency increased. Adaxial stomatal traits (density, ratio, and occurrence) had nonlinear relationships with ancestry, with higher adaxial stomatal density and occurrence at intermediate ancestry, but hybrids with high *P. trichocarpa* ancestry showing adaxial stomatal traits more like *P. trichocarpa* (Figure 2A–3A).

**Figure 2.**
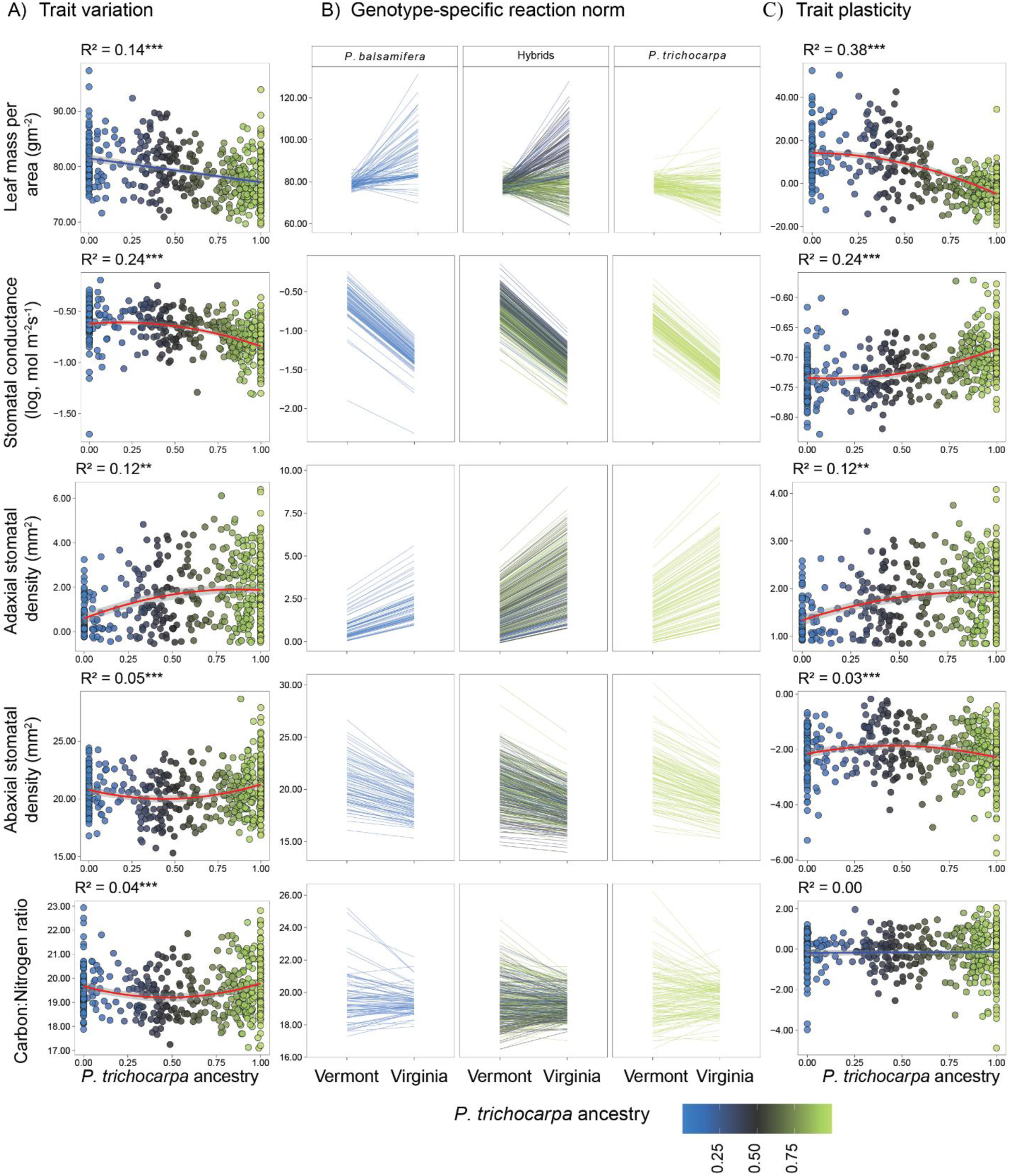
(A) Relationship between genomic ancestry and trait variation, estimated as genotype-level BLUPs. (B) Reaction norms for physiological traits across common gardens. Lines represent genotype-specific reaction norms estimated from linear mixed models. Genotypes were grouped into genotypic classes for visualization only. (C) Relationship between genomic ancestry and trait plasticity, quantified as the slope of the reaction norm. Blue and red lines represent the best-fit linear and quadratic models, respectively, and the shaded gray area indicates the 95% confidence interval. The coefficient of determination (R^2^) for the selected model is shown in each panel, with asterisks indicating statistical significance (*p* < 0.05*, *p* < 0.01**, *p* < 0.001***).

**Figure 3.**
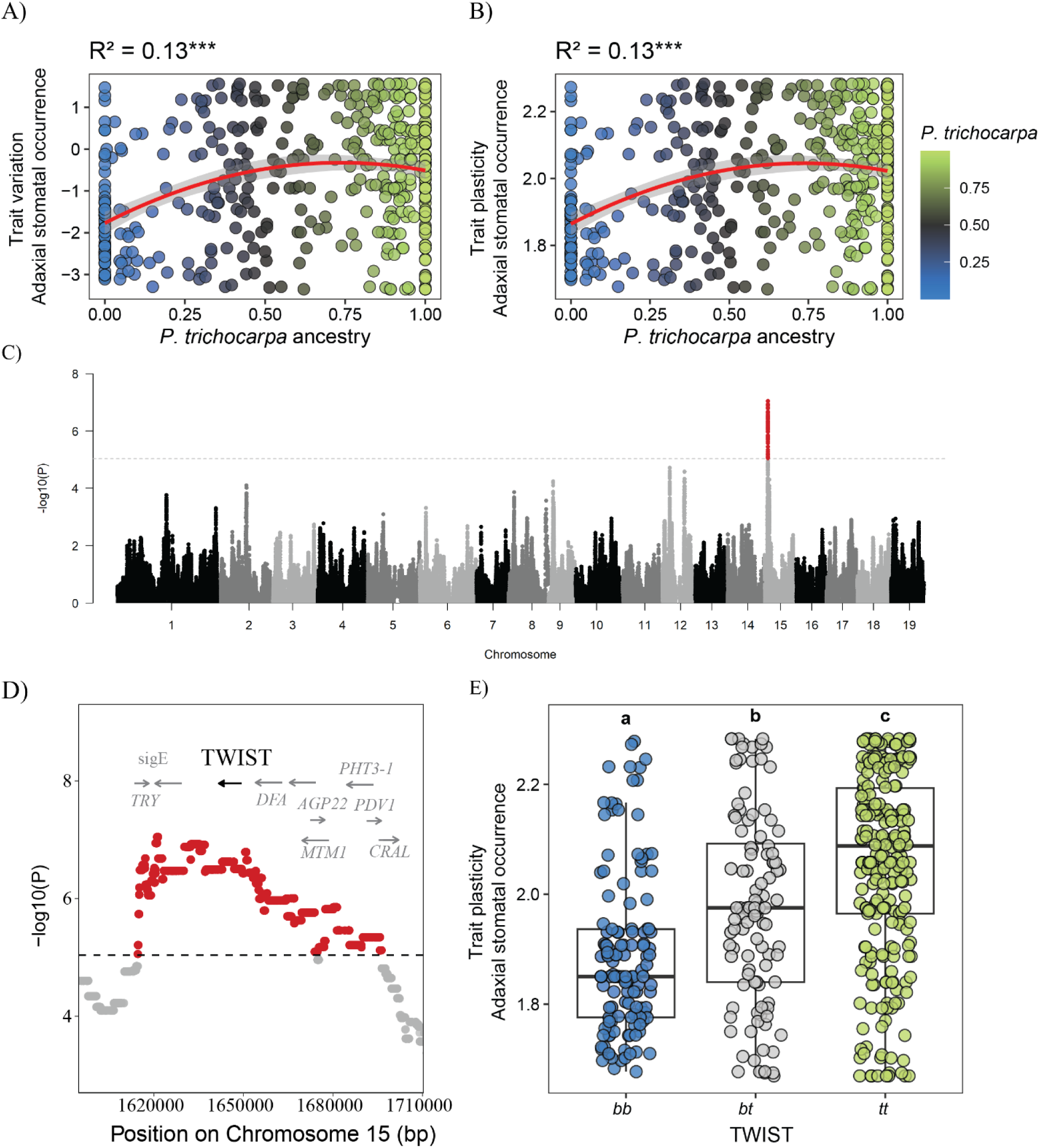
(A) Relationship between ancestry and adaxial stomatal occurrence variation, estimated as genotype-level BLUPs. (B) Relationship between genomic ancestry and plasticity in adaxial stomatal occurrence, quantified as the slope of the reaction norm. Red lines show best-fit quadratic models, with shaded gray areas indicating 95% confidence intervals. R² values are shown, with asterisks indicating statistical significance (p < 0.001 ***). (C) Manhattan plot showing results from GWAS by admixture mapping for plasticity in adaxial stomatal occurrence. The black dashed line indicates the genome-wide significance threshold (–log₁₀ (P) ≈ 5.004), corrected for admixture burden (0.05/5055). (D) Zoomed-in view of the candidate region on chromosome 15 showing genes identified. (E) Boxplots display variation in plasticity for adaxial stomatal occurrence across local ancestry classes at *TWIST* (homozygous *P. balsamifera* (*bb*), heterozygous (*bt*), and homozygous *P. trichocarpa* (*tt*) genotypes). Positive slope values indicate an increase in adaxial stomatal occurrence across environments, whereas negative values indicate a decrease. The dashed gray line represents no plasticity. Letters indicate results from Tukey’s post hoc test following ANOVA; different letters denote statistically significant differences among genotypic classes.

Similar with trait variation, genomic ancestry influenced the magnitude and direction of reaction norms across trait classes (Figure 2B-C). Genotypes with higher *P. trichocarpa* exhibited greater plasticity for photochemical (*p* > 0.05, Figure S4, Table S2), and adaxial stomatal traits (*p* > 0.01). In contrast, genotypes with higher *P. balsamifera* ancestry exhibited greater plasticity for leaf dry mass (*slope* = -0.28 g, *p* > 0.01), leaf area (*slope* = 0.20 m^2^, *p* > 0.001), leaf mass per area (*slope* = -39.59 gm^-2^, *p* > 0.001), and stomatal conductance (*slope* = 0.14 mol m^−2^ s^−1^, *p* > 0.001). Hybrids had intermediate plasticity for leaf area, and photochemical traits. However, for some traits, the relationship between ancestry and plasticity was nonlinear (Table S2), with hybrids exhibiting transgressive reaction norms or resembling one parental species more closely. Specifically, hybrids showed reduced plasticity for leaf dry mass and abaxial stomatal density with respect to parental genotypes, but higher plasticity in stomatal conductance. For plasticity in adaxial stomatal traits, hybrids exhibited greater plasticity than *P. balsamifera* genotypes, with plasticity steeply increasing for genotypes with intermediate ancestry and remaining similarly higher with *P. trichocarpa* ancestry (Figure 2C–3B). In contrast, leaf dry mass plasticity exhibited a nonlinear relationship with ancestry, with increased plasticity for hybrids and reduced plasticity with greater *P. trichocarpa* ancestry (Figure 2C). Genomic ancestry did not significantly (*p* > 0.05) influence plasticity for water-use efficiency, adaxial and abaxial pore length, nitrogen content, nitrogen isotope composition, or carbon-to-nitrogen ratio, suggesting that plasticity in these traits is likely independent of genomic ancestry.

### Common genetic basis for trait variation and their environmental responsiveness identified via admixture mapping

Admixture mapping for trait variation and trait plasticity (slopes of the reaction norm) identified 11 genes that had a shared genetic basis underlying both adaxial stomatal trait differences and their plasticity (Figure 3C, Figure S5-S6, Table S3). Within a region of ∼ 0.23 Mb on chromosome 15, ten genes including transcription factor *TWIST* (*Potri.015G022300*) were associated with adaxial stomatal occurrence variation and its plasticity, suggesting a role in both trait expression and its regulation in response to the environment (Figure 3D). Comparison across local ancestry classes revealed that homozygous *P. balsamifera* genotypes exhibited reduced plasticity for adaxial stomatal occurrence relative to homozygous *P. trichocarpa*, with heterozygotes exhibiting intermediate values (Figure 3E).

Beyond chromosome 15, six additional genes were associated with reaction norms, indicating genetic control of environmental responsiveness (Table S3). These include three candidate genes associated with plasticity for stomatal density ratio, including a DHHC palmitoyltransferase (*Potri.005G104700*), and a Casparian strip membrane protein (*Potri.009G014600*), in chromosomes 05 and 09, respectively. Plasticity for stomatal conductance was associated with three genes in chromosome 10: two phenylalanine ammonia-lyase genes (*PAL2, Potri.010G224100* and *Potri.010G224200*) and an adenosine kinase (*ADK, Potri.010G224300*). In contrast, for leaf morphological traits no genes were significantly associated with plasticity. Instead, several loci were linked exclusively to trait differences in leaf area and leaf dry mass, including two membrane-associated kinase regulators (MAKR; *Potri.017G115700*, *Potri.017G115801*) and a leucine-rich repeat protein kinase (*Potri.017G115900*) located on chromosome 17.

To account for the high correlation between traits and their plasticity (Figure S2), we repeated admixture mapping using scaled plasticity estimates. Under this scaled analysis, no loci were significantly associated with plasticity. This suggests that the associations detected may reflect genetic effects on trait values, or pleiotropic loci influencing both trait expression and plastic responsiveness.

### Climate warming predicts directional changes in genetic variation underlying trait and trait plasticity

Comparisons of historical climate with future climate projections at genotype collection sites in the hybrid zone predict substantial shifts across multiple climatic variables, with higher-emission scenarios having larger differences from historical climates (Figure 4). Across all SSPs, mean annual temperature, degree-days above 5 °C, extreme minimum temperature, and the length of the frost-free period (FFP, NFFD, EMT, eFFP) are projected to increase relative to the historical climate, while chilling degree-days (DD_0) decline indicating substantial warming. Moisture indices (e.g., CMD, CMI, Eref) further suggest progressive drying, particularly under SSP370 and SSP585, whereas precipitation variables exhibit more variable responses across projected scenarios.

**Figure 4.**
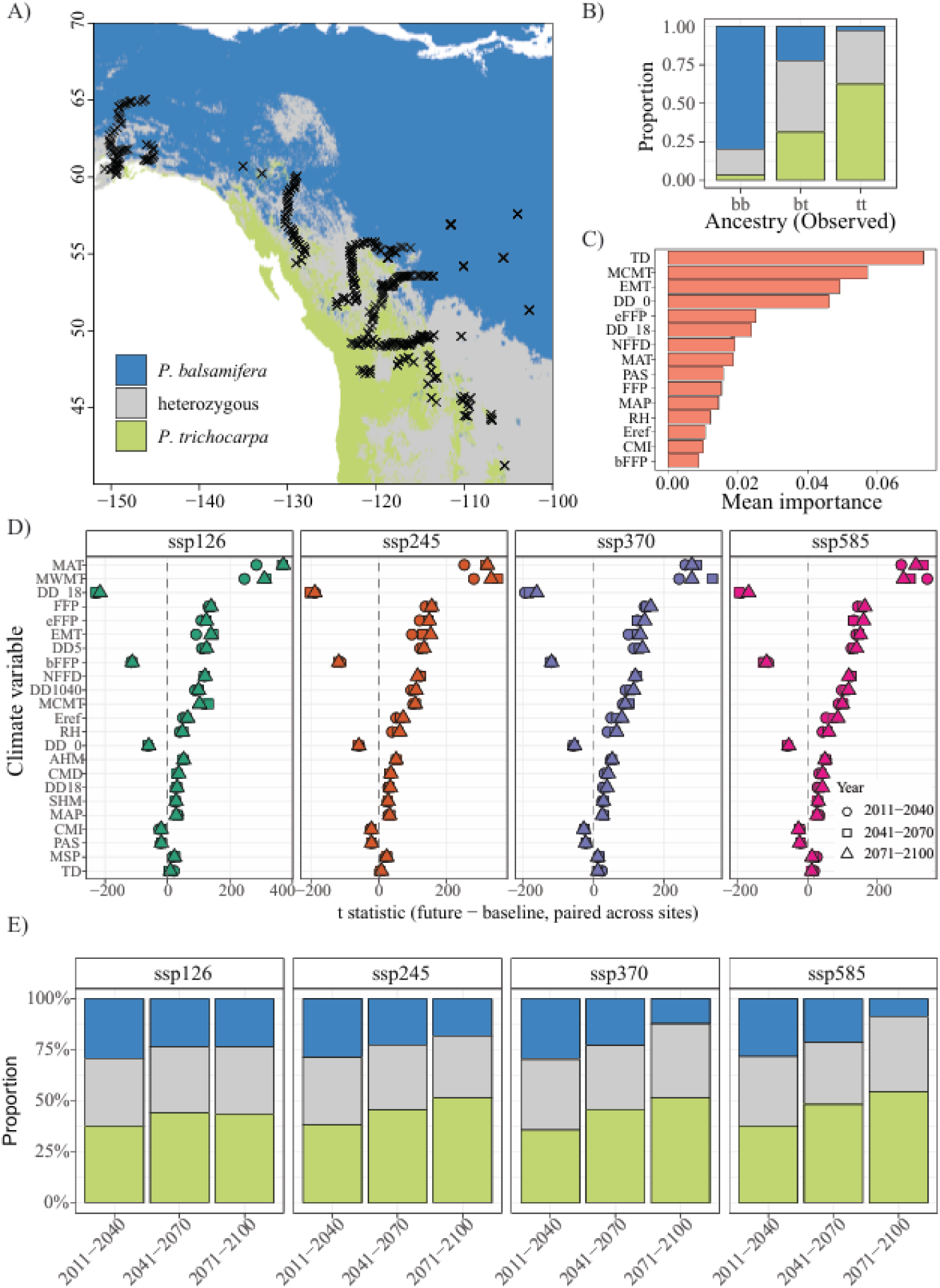
Analysis of *TWIST* associated with adaxial stomatal traits in the hybrid zone. (A) Geographic distribution of predicted ancestry classes at genotype sites in the hybrid zone in the baseline climates. (B) Cross-validation confusion matrix showing the proportion of correctly and incorrectly classified ancestry classes. (C) Top 15 climate variables ranked by mean importance across 100 random forest iterations. (D) Projected changes in climate variables at genotype sites relative to the historical baseline under four Shared Socioeconomic Pathways (SSPs) and three future periods (2011–2040, 2041–2070, 2071–2100). (E) Predicted distribution of ancestry classes at genotype sites across SSPs and time periods across 100 random forest replicates.

We trained random forest models using historical climatic data at genotype sampling sites to quantify associations between climate and local ancestry at *TWIST* (Figure 4A). This gene was selected because local ancestry at this locus was associated with both adaxial stomatal density and occurrence differences as well as their plasticity. Model evaluation indicated that the trained random forest models captured associations between historical climate and local ancestry with moderate accuracy (mean accuracy = 0.64 ± 0.03, κ = 0.45 ± 0.04 across 100 iterations). Misclassification of local ancestry was most frequent for heterozygous genotypes, likely reflecting their intermediate position between homozygous genotypes (Figure 4B). Random forest models identified temperature-related variables as the strongest predictors of local ancestry at *TWIST* (Figure 4C). The most influential variables were continentality (TD), mean coldest month temperature (MCMT), extreme minimum temperature (EMT), and chilling degree-days (DD_0). These results emphasize that local ancestry is associated with seasonal climatic variables, which are projected to shift under future climates (Figure 4D).

Climate projections predicted substantial shifts in the distribution of local ancestry in *TWIST* under climate change scenarios (Figure 4E). Across all scenarios, the frequency of *P. trichocarpa* ancestry (*tt*) was predicted to increase, particularly under higher-emission pathways (SSP370, SSP585). In contrast, *P. balsamifera* ancestry (*bb*) was predicted to decline, while heterozygous genotypes (*bt*) remained constant. To account for stochasticity introduced by down-sampling, projections were generated from 100 independently resampled model runs. Variation among iterations was small relative to the magnitude of the predicted shifts (SD = 0.20 – 0.29), providing strong support for projected increases in *P. trichocarpa* ancestry and declines in *P. balsamifera*, with heterozygotes relatively constant over time under all scenarios.

## Discussion

Identifying the genetic variation underlying plasticity for traits that regulate phenotypic expression across environments is essential for predicting population response to climate change. By integrating phenotypic, genomic, and climate of origin data for genotypes grown in common gardens originating from the *Populus trichocarpa × P. balsamifera* hybrid zone, we identified the genetic variation contributing to key traits that regulate plants response to environmental change. Specifically, we found substantial genetic variation for plasticity within the hybrid zone, as G×E explained a considerable proportion of phenotypic variance across trait classes. While hybrid genotypes in general exhibited intermediate plasticity relative to parental genotypes, for stomatal traits they exhibited transgressive reaction norms. Using GWAS by admixture mapping, we identified regulatory hotspots underlying adaxial stomatal traits and their plasticity, including *TWIST*, suggesting that their evolution may not be independent. Climatic projections suggest that shifts in climate are likely to favor *P. trichocarpa* alleles associated with stomatal trait regulation and plasticity across the hybrid zone, while *P. balsamifera* alleles may persist in hybrids through heterozygosity. Overall, this study shows that hybridization may influence genetic variation underlying plasticity impacting genotypes’ response to the environment, and that genetic variation maintained within hybrids will be critical component of adaptation in response to future climates.

### Genetic variation for plasticity in physiological traits exists in the hybrid zone

A substantial proportion of trait variance was explained by G×E interactions, indicating that genetic variation for plasticity exists for genotypes within the *P. trichocarpa* × *P. balsamifera* hybrid zone. Previous studies within this hybrid zone suggested that genotypes with differing genomic ancestries exhibited strong G×E effects for fitness and physiological traits when grown in warmer environments (Mead et al. 2026a,b). Consistent with this, we found that G×E explained a substantial proportion of variation across trait classes, with particularly strong effects for leaf mass per area and nitrogen content (9.86–35.16%), indicating potential for these traits to evolve in response to environmental change. Given that leaf mass per area and nitrogen content are associated with structural investment and photosynthetic capacity, genetic variation for plasticity suggests that, under varying environmental conditions, selection may shape genotypes’ ability to adjust carbon gain and resource-use strategies (Wright et al. 2004; John et al. 2017; Onoda et al. 2017; Hu et al. 2025). Stomatal traits had a high proportion of variance explained by genetic effects (G), with smaller but significant G×E contributions (5.29–7.64%), indicating that most variation reflects the evolution of differences among genotypes, though there is a genetic component to the regulation of phenotypic expression. Regardless of whether selection acts on trait variation, plasticity, or both, genetic variation in stomatal traits and their plasticity provides the raw material for natural selection, demonstrating the evolutionary potential of traits and their plastic responses under changing environmental conditions.

Despite substantial contributions of genotype, environment, and G×E, a large portion of trait variation remained unexplained. This likely reflects biological complexity not captured by the model, including microclimatic heterogeneity and temporal variability (Westneat et al. 2015). Given that our data comes from two common gardens sampled in one year, unmodeled spatial and temporal stochasticity likely contributed to residual variance, underscoring the need for temporal replication and replication of gardens across environmental gradients.

### Genomic ancestry shapes trait expression and the magnitude and direction of reaction norms across trait classes

Genomic ancestry was associated with both trait variation and plasticity for genotypes within the hybrid zone, suggesting that interspecific gene flow can be an important contributor to trait differences and their regulation across environments. For trait variation, we found that genotypes with higher *P. trichocarpa* ancestry exhibited greater leaf dry mass, leaf area, intrinsic water-use efficiency, and adaxial stomatal traits, whereas *P. balsamifera* genotypes had higher leaf mass per area, stomatal conductance, and photochemical efficiency (Figure 2, S3). These patterns suggest climate adaptation differentiates the parental species, with *P. trichocarpa* exhibiting a more acquisitive strategy associated with higher carbon gain, and *P. balsamifera* a more conservative strategy that may confer greater resilience to stresses, such as water limitation. Genomic ancestry also influenced both the magnitude and direction of reaction norms across trait classes. In our study, *P. trichocarpa* exhibited greater plasticity than *P. balsamifera* in adaxial stomatal traits. Increased adaxial stomatal density can contribute to CO₂ uptake and transpirational cooling in response to warmer conditions (Drake et al. 2019; Muir 2019; Wall et al. 2022), but can also be associated with potential tradeoffs with increased pathogen susceptibility to poplar leaf rust, *Melampsora* spp. (Fetter et al. 2021; Fetter and Keller 2023). Yet, *P. balsamifera* showed higher plasticity in leaf mass per area, leaf area, and leaf mass than *P. trichocarpa*, indicating a greater capacity of these genotypes to modify leaf tissue investment in response to changing environments. These contrasting patterns in plasticity among traits suggest that the parental species differ in physiological strategies, potentially reflecting adaptation to their climates of origin (Eisenring et al. 2022; Solé-Medina et al. 2022; Ramírez-Valiente et al. 2025; Mead et al. 2026b).

Recombination between the divergent genetic architectures resulted in a range of plastic responses. For some traits, hybrids exhibited greater plasticity (steeper slopes) than parental genotypes, whereas for others their reaction norms resembled one parental species (Figure 2C, S4, Table S2). Specifically, our models indicated that hybrids exhibited greater plasticity for stomatal conductance, but lower plasticity for abaxial stomatal density relative to parental species. For adaxial stomatal traits, hybrids displayed reaction norms similar to those of *P. trichocarpa*. Higher plasticity in adaxial stomatal traits and stomatal conductance may enhance carbon uptake under warm or high-light conditions by increasing stomatal abundance on the adaxial leaf surface (Haworth et al. 2018; Drake et al. 2019; Harrison et al. 2020; Wall et al. 2022). However, these responses may also increase water loss and susceptibility to foliar pathogens (McKown et al. 2014; Fetter et al. 2021; Fetter and Keller 2023). In contrast, abaxial stomatal density in hybrids exhibited reduced plasticity, with genotypes generally having fewer abaxial stomata but denser adaxial stomata from Vermont to Virginia environments (Figure 2B). This pattern suggests a redistribution of stomatal investment between leaf surfaces, potentially optimizing gas exchange while maintaining water balance under the warmer and drier conditions of Virginia compared with Vermont. Notably, we found no relationship between ancestry and plasticity for water-use efficiency. These suggest that hybrid genotypes compensate for higher stomatal density by reducing stomatal conductance, thereby maintaining similar gas exchange per unit leaf area. Thus, while hybridization can create novel allelic combinations that may facilitate the evolution of plasticity (Schwartz et al. 2024), the adaptive value of these responses ultimately depends on the balance between carbon gain and water conservation across environments and their effects on fitness (Mead et al., 2026).

### Shared genetic architecture constrains the evolution of plasticity in stomatal traits

Previous studies report no association between trait variation and plasticity, suggesting independent genetic architectures (Jin et al. 2023; Love and Ferris 2024). In contrast, our results show strong correlations between traits and their plasticity and identify candidate genes associated with both, consistent with a shared genomic architecture underlying trait expression and its regulation. Specifically, in this study, chromosome 15 emerged as a hotspot for associates between adaxial stomatal variation and its plasticity, with *TWIST* and nine additional genes consistently associated with both. This shared genetic architecture may constrain the independent evolution of plasticity (Via 1993; Lafuente et al. 2024), particularly if future environments require changes in environmental responsiveness that are not aligned with the direction of selection on adaxial stomatal variation. *TWIST*, the *Populus* homolog of *Arabidopsis SPEECHLESS* (*SPCH*), regulates stomatal lineage initiation, and its expression is influenced by environmental cues (Lau et al. 2018) including high temperatures and drought (Viger et al. 2016; Lau et al. 2018). Transcriptomic data for *P. trichocarpa* indicates that *TWIST* expression is developmentally regulated, with highest expression in actively developing leaf tissue followed by a decline as leaves mature (Goodstein et al. 2012; McKown et al. 2019). For genotypes collected in this study, functional validation will be needed to determine whether environmentally induced changes in *TWIST* expression translate into shifts in stomatal patterning.

In this study, *P. balsamifera* genotypes that were homozygous for *TWIST* exhibited reduced plasticity for adaxial stomatal occurrence, while *P. trichocarpa* exhibited greater plasticity (Figure 3E). Heterozygotes exhibited intermediate plasticity consistent with an additive effect of allelic ancestry at this locus. Adaxial stomatal occurrence increased from Vermont to Virginia for all genotypes, indicating a directional response to the change in environment. At the same time, the slope of the reaction norms differed among local ancestry classes at *TWIST*, suggesting sensitivity in the magnitude of ancestry-specific allele effects varies across environments (Des Marais and Juenger 2010). Together, these findings suggest that *TWIST* plays a key role in regulating adaxial stomatal development in response to environmental change for *Populus*.

Genetic correlations suggest there can be a shared genetic architecture influencing traits and their expression (Scheiner 1993; Pigliucci 2005). Alternatively, genotypes with higher trait means may appear more plastic because larger trait values can show greater absolute differences between environments due to scaling effects. These absolute differences may not reflect intrinsic variation in environmental responsiveness among genotypes. To account for this, we scaled plasticity relative to genotype-specific intercepts to control for differences in trait means and performed GWAS by admixture mapping using scaled plasticity. However, no significant associations were detected. This suggests that loci identified using unscaled plasticity primarily reflect shared genetic control of adaxial stomatal trait variation and plasticity, rather than loci affecting plasticity independently of trait variation.

### Future climatic conditions are predicted to favor *P. trichocarpa* and heterozygous genotypes at *TWIST*

Climate change is expected to modify species distributions by altering extrinsic selective pressures (Taylor et al. 2015). Our results indicate that the climatic conditions favoring different ancestry at key regulatory genes such as *TWIST* may shift, with *P. trichocarpa* alleles favored under future climates within the *P. trichocarpa* × *P. balsamifera* hybrid zone (Figure 4). Our random forest models identified temperature-seasonality variables, including continentality, mean coldest month temperature, extreme minimum temperature, and chilling degree-days, as the strongest predictors of local ancestry at *TWIST*, suggesting that temperature extremes have exerted selection pressure on local ancestry for this candidate gene. These predictors align with the expectations for parental species, with *P. balsamifera* adapted to continental climates and *P. trichocarpa* to more maritime conditions (Richardson et al. 2014; Suarez-Gonzalez et al. 2018; Zavala-Paez et al. 2025). Climate scenarios from climateNA, which project warming and reduced winter severity, forecast an expansion of *P. trichocarpa* ancestry at *TWIST* for all SSPs. Across our common gardens, homozygous genotypes for *P. trichocarpa* alleles at *TWIST* showed higher adaxial stomatal density and plasticity, with leaves developing more adaxial stomata in the warmer Virginia common garden. This increase in adaxial stomatal density and occurrence may alter key physiological trade-offs. Increased density of adaxial stomata could enhance transpirational cooling and CO₂ uptake under projected warmer conditions, but within increasingly arid regions of the hybrid zone, this same response may elevate the risk of water loss through transpiration (Mott et al. 1982; Drake et al. 2019). These opposing outcomes suggest that adaptive responses to warming may be constrained by trade-offs between carbon gain and water conservation. Although *P. balsamifera* homozygotes at *TWIST* are predicted to decline in response to climate change, the *P. balsamifera* allele is expected to persist in heterozygotes. These suggest that hybridization may provide an important mechanism to maintain allelic diversity across systems. Genotypes with heterozygous alleles for *TWIST* exhibit intermediate plasticity for adaxial stomatal traits relative to the parental species, potentially reflecting adaptation to intermediate environments. Our findings align with Mead et al. (2026a,b) who forecast that genotypes with higher *P. trichocarpa* ancestry are likely to outperform *P. balsamifera* when considering genome-wide ancestry. Together, these results suggest that the pattern observed at *TWIST* is consistent with broader selection favoring *P. trichocarpa* ancestry, while also highlighting *TWIST* as a candidate locus regulating adaptive responses.

Although our models forecast an increase in *P. trichocarpa* ancestry at *TWIST*, these projections assume that *Populus* populations can evolve or migrate at a rate comparable to the pace of climate change (Aguirre-Liguori et al. 2021). Standing genetic variation and introgression, particularly through maintenance of heterozygosity at *TWIST,* likely represent the primary sources of adaptive variation over contemporary timescales. While *Populus* disperses pollen and seeds effectively through wind and water (Slavov et al. 2009), migration is often constrained by geographic barriers such as the Rocky Mountains, which can limit the ability of genotypes to track rapidly shifting climates (Aitken et al. 2008; Bolte et al. 2024; Mead et al. 2026a). Moreover, climate-driven changes in co-occurring species are likely to alter community composition and biotic interactions (Taylor et al. 2015; Liang et al. 2018). These ecological and evolutionary factors add further uncertainty to our forecasts and emphasize the need to integrate dispersal limits, landscape structure, and interspecific interactions into our models. Whether selection can act rapidly enough to shift the geographic distribution of these alleles remains unclear. However, genetic variation maintained within the hybrid zone may expand the geographic extent over which beneficial alleles are present, potentially increasing the capacity for adaptive responses under future climates.

## Conclusions

Our study demonstrates that hybridization can be an important source of genetic variation that translates into diverse reaction norms across environments for *Populus* genotypes, extending the phenotypic range of responses beyond those expressed by either parental species. A shared genetic basis for trait variation and its plasticity suggests that maintaining this genetic variation is essential, as it underlies both trait differences and environmental responsiveness. However, shared genetic architecture may constrain adaptation if plasticity cannot evolve independently of trait variation. We predict that changing environmental conditions will select for genetic variation underlying plasticity for future climates. If trees can migrate or persist within suitable regions, *P. trichocarpa* genotypes are expected to increase in frequency; however, hybrids may preserve allelic diversity in the heterozygous form, maintaining the genetic diversity necessary for future adaptation. Taken together, these findings highlight that hybridization not only influences plastic responses but also contributes to the long-term evolutionary potential of forest trees facing rapid climatic change.

## Data availability

The data, code and custom scripts for all analyses are available on GitHub: https://github.com/michestzav/Genetic_Variation_Plasticity.git. Scripts for local ancestry inference and GWAS by admixture mapping can be found here: https://github.com/michestzav/Stomata_evolution_Populus.git

## Author contributions

Michelle Zavala-Paez led the study design, data collection, data analysis, interpretation of results, and manuscript preparation. Alayna Mead contributed to data analysis, interpretation, and writing of the manuscript. Baxter Worthing and Sara Klopf assisted with data collection and contributed to manuscript writing. Stephen Keller contributed to data collection, data analysis, and manuscript preparation. Jason Holliday and Matthew C. Fitzpatrick contributed to data collection and manuscript writing. Jill Hamilton contributed to study design, data collection, data analysis, interpretation of results, and manuscript writing.

## Funding

This research was supported by the National Science Foundation grant PGR-1856450, the USDA National Institute of Food and Agriculture (NIFA) project and Hatch Appropriations (PEN04809, Accession 7003639), and NIFA project VA-136641. Additional support was provided by the Schatz Center for Tree Molecular Genetics, the Huck Institutes of the Life Sciences, the Department of Ecosystem Science and Management, and the Graduate Program in Ecology at Pennsylvania State University.

## Conflict of interest

The authors declare no conflicts of interest

## Acknowledgments

We are grateful to Tommy Phannareth for his support during physiological trait sampling in the Virginia common garden. We also acknowledge the assistance of Kyle Peer, Clay Sawyers, and Deborah Bird at the Virginia Tech Reynolds Homestead Forestry Research Station for their role in plant propagation. Nadia Garzione contributed to the collection and processing of physiological trait data, and we thank her for her efforts. We further appreciate the members of the Hamilton Lab, Kyra LoPiccolo, Diego del Orbe, Muqing Liu, Mary McCafferty, and Sammy Muraguri, for their thoughtful feedback and discussions during lab meetings, which contributed to the development of this study.

## Notes

### Competing Interest Statement

The authors have declared no competing interest.

