## Supplementary_material for "Genetic variation underlying plasticity in physiological traits mediates response to climate in a *Populus* hybrid zone"

### Contents

Supplementary Methods

Figure S1. Variance partitioning of physiological traits, showing contributions of genotype, environment, G×E, block, and residual variance
Figure S2. Spearman's rank correlations between trait variation and trait plasticity
Figure S3. Relationship between *P. trichocarpa* ancestry and trait variation estimated as best linear unbiased predictors (BLUPs) across genotypes
Figure S4. Relationship between *P. trichocarpa ancestry* and trait plasticity, quantified as genotype-specific reaction norm slopes across genotypes.
Figure S5. Quantile–quantile (QQ) plots of association results for trait variation in *Populus* hybrids
Figure S6. Quantile–quantile (QQ) plots of association results for slopes of reaction norms in *Populus* hybrids

Table S1. Associations between genomic ancestry and trait variation, including model selection, fit ($R^2$), AIC, and parameter estimates.
Table S2. Associations between genomic ancestry and trait plasticity, including model selection, fit ($R^2$), AIC, and parameter estimates.
Table S3. List of candidate genes underlying physiological trait variation and plasticity identified through admixture mapping analysis.

### Supplementary methods

#### Library preparation, sequencing and variant calling

Genomic libraries were sequenced on an Illumina NovaSeq 6000 using an S4 flow cell in 2 × 150 bp paired-end format, with 64 samples per lane. Illumina reads for each genotype were aligned to the *Populus trichocarpa* reference genome (v4.0), generating SAM files that were subsequently converted to BAM format using SAMtools (Li *et al.*, 2009). Individual gVCF files were generated using the HaplotypeCaller algorithm in GATK v3.7 and subsequently combined into a single VCF file using the GenotypeGVCFs function. The initial dataset included ~82 million variants and was filtered based on standard quality metrics. Filters included mapping quality (MQ < 40.00), strand bias (FS > 40.000, SOR > 3.0), mapping quality rank sum (MQRankSum < –12.500), read position bias (ReadPosRankSum < –8.000), and depth of coverage (QD < 2.0). INDELs and SNPs with more than two alternate alleles were excluded, resulting in a final dataset of 29,663,130 high-quality biallelic SNPs. Further filtering removed SNPs with a minor allele frequency below 5% and more than 10% missing data, yielding a final dataset of 7,167,726 biallelic SNPs for downstream analyses.

#### Phenotyping

For each genotype, gas exchange, light-use efficiency and stomatal traits were assessed using the first fully expanded leaf. This leaf was selected to minimize age and environmental effects between samples (Fetter *et al.*, 2021). Gas exchange and light efficiency traits including stomatal conductance (gsw, mmol $m^{-2}$ $s^{-1}$), electron transport rate (ETR, mol $m^{-2}$ $s^{-1}$), quantum efficiency of PSII (ΦPSII), minimum (Fs), and maximum fluorescence in light (Fm), were measured using a LI-600 porometer in three non-overlapping areas per each fully expanded leaf without detaching the leaf from the plant. Thus, individual ΦPSII, ETR, Fs, Fm and $g_{sw}$ values represent the average of these three measurements.

After measuring these traits, leaves were collected to evaluate stomatal density, size, and distribution. To capture variation across both leaf surfaces, a thin layer of Newskin liquid bandage was applied to the adaxial and abaxial leaf surface, creating two impressions per individual. Impressions were mounted on microscope slides, and images were taken using an Olympus BX-53 microscope paired with a DP23 digital camera. All micrographs were scaled to a 0.37 × 0.25 mm grid for consistency. Stomatal size traits were analyzed separately for each surface using ImageJ (Abràmoff *et al.*, 2004), focusing on pore length of adaxial ($P_U$, μm) and abaxial ($P_L$, μm) stomata. For each image, four evenly spaced lines were drawn, and five stomata were measured to calculate average values per surface. For stomatal density, adaxial ($D_U$, $mm^{-2}$) and abaxial ($D_L$, $mm^{-2}$) density were estimated using LeafNet (Li *et al.*, 2022), with density defined as the number of stomata per square millimeter of leaf area. To validate the accuracy of automated estimates, stomatal density was manually counted from micrographs of 100 randomly selected genotypes. Manual and automated counts were strongly correlated ($r = 0.95$, $p < 0.05$; Figure S1), supporting the reliability of the automated method, therefore automatic counts were used for all genotypes in the study. To characterize overall stomatal abundance, total stomatal density ($D_T$) was calculated by summing the adaxial ($D_U$) and abaxial ($D_L$) stomatal densities, providing a measure of the total number of stomata per unit leaf area. The degree of amphistomy was quantified using the stomatal ratio (SR), defined as the proportion of adaxial stomata relative to the total stomatal count ($SR = D_U / D_T$). This ratio ranges from 0 (stomata present only on the abaxial surface) to 1 (stomata present only on the adaxial surface). To further assess variation in adaxial stomatal placement, we recorded the presence (1) or absence (0) of stomata on the adaxial surface, referred to as adaxial stomatal occurrence ($O_U$).

Using the second fully expanded leaf, we measured leaf structure, size and isotopic composition. To estimate leaf structure and size, each leaf was scanned with an Epson Perfection V39 scanner to generate a high-resolution digital image. Leaf area ($m^2$) was then quantified using ImageJ software (Abràmoff *et al.*, 2004). Following scanning, each leaf was dried at 60 °C until a constant weight was reached. The dry mass (g) was then recorded. Leaf mass per area (LMA) was calculated by dividing the dry mass by the corresponding leaf area ($gm^{-2}$). To estimate elemental and isotopic composition, dried leaves were homogenized using a TissueLyser II (Qiagen, Germany). Approximately 2 mg of ground tissue per sample was weighed into tin capsules and analyzed for carbon (C, %), nitrogen (N, %), and stable isotope of carbon ($\delta^{13}C$, ‰) and nitrogen ($\delta^{15}N$, ‰) at the Central Appalachians Stable Isotope Facility (CASIF), University of Maryland Center for Environmental Science (USA).

**Admixture mapping to identify the genomic basis of physiological trait variation and their plasticity**

To associate locus-specific ancestry with trait variation and trait plasticity, we applied separate univariate linear mixed models for each trait and its plasticity using GEMMA v0.94.1 (Zhou & Stephens, 2014). Models included a kinship matrix (computed with the -gk 1 option) to correct for genetic relatedness, and ancestry ($K = 2$) as a covariate to account for background population structure (Fetter & Keller, 2023). To correct for multiple testing, we applied a significance threshold based on the admixture burden, which estimates the number of effectively independent chromosomal segments in the genome (Shriner *et al.*, 2011). Admixture burden was calculated by fitting an autoregressive model to local ancestry tracts in each genotype and estimating spectral density at frequency zero using the *spectrum0.ar* function from the *coda* package in R (Plummer *et al.*, 2006). We then averaged the number of independent ancestry blocks across individuals to obtain the effective number of tests resulting in a mean of 5,055 independent tests (Shriner *et al.*, 2011; Fetter & Keller, 2023). Accordingly, the admixture burden threshold was set to 5.004, calculated as $-\log_{10}$ (0.05/5,055). Manhattan plots were generated using the *qqman* package in R (Turner, 2018) to visualize the results, with significant loci identified according to the admixture burden-adjusted threshold. QQ plots were used to verify that population structure was controlled adequately.

### Supplementary figures

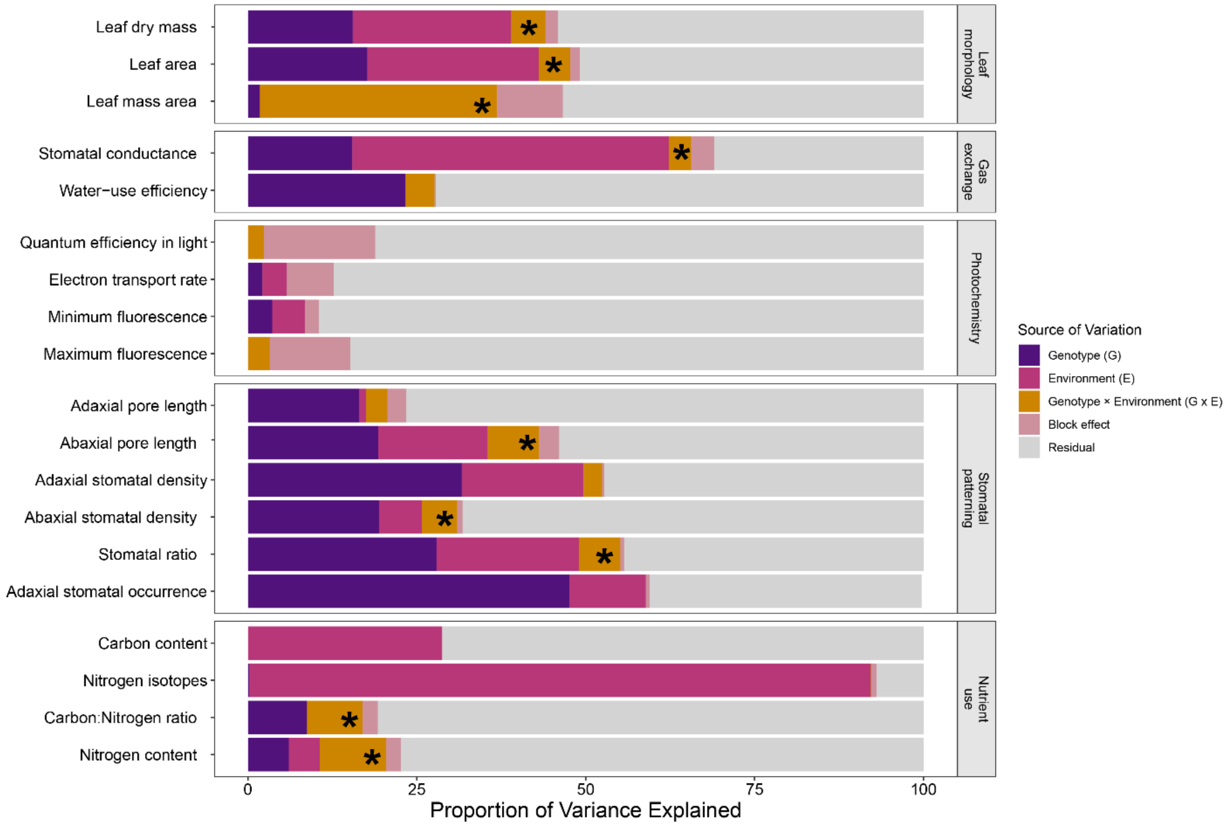


**Figure S1**. Bar plots show the proportion of variance explained by genotype (purple), environment (pink), genotype × environment interaction (orange), block effect (light pink), and residual variance (gray) for each physiological trait. Traits with statistically significant genotype × environment (G×E) interactions, based on likelihood ratio tests (LRT) comparing models with and without the interaction term, are marked with an asterisk.

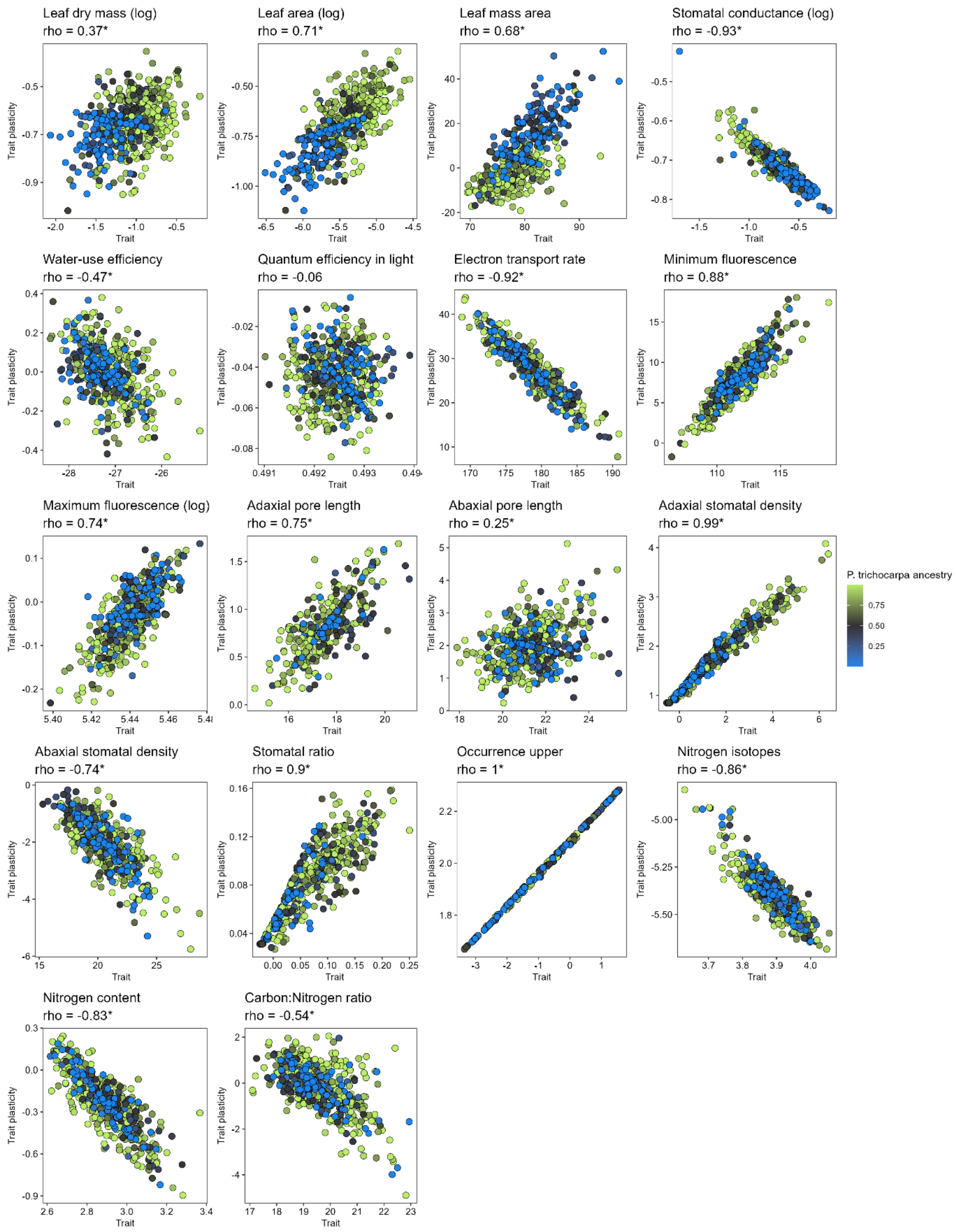


**Figure S2.** Spearman's rank correlations (ρ) between trait variation (x-axis) and trait plasticity (y-axis). Statistically significant correlations are denoted by an asterisk (*, $p < 0.05$).

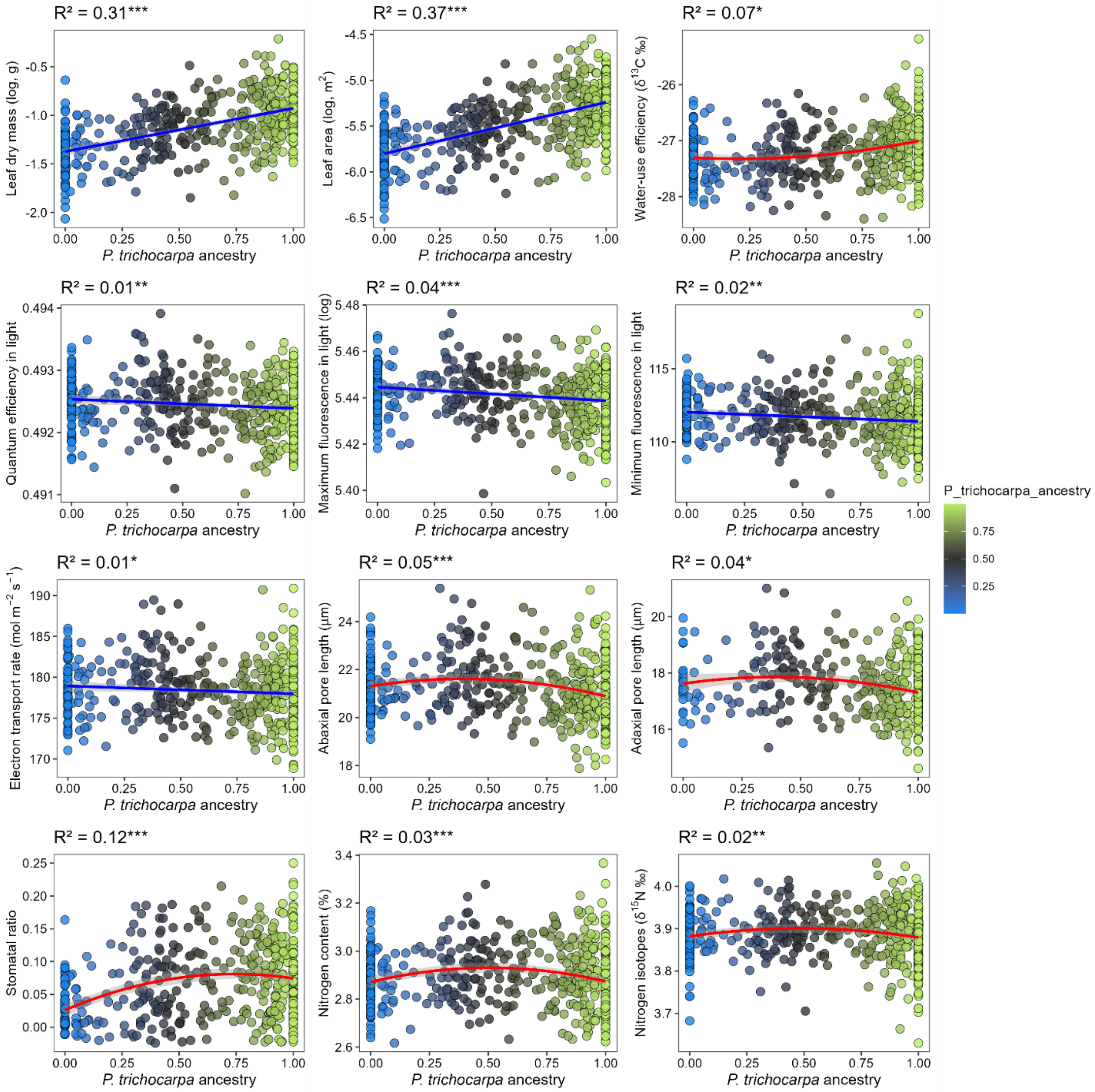


**Figure S3.** Relationship between *P. trichocarpa* ancestry (x-axis) and trait values estimated as best linear unbiased predictors (BLUPs; y-axis) across genotypes. Points are colored by genomic ancestry, with values near 0 indicating greater *P. balsamifera* ancestry (blue), values near 1 indicating greater *P. trichocarpa* ancestry (green), and intermediate values representing admixed genotypes (gray). Solid lines represent the best-supported model for each trait, selected using AIC comparison between linear and quadratic models ($\Delta AIC > 2$). Linear fits are shown in blue and quadratic fits in red, with shaded areas representing 95% confidence intervals. The coefficient of determination ($R^2$) for the selected model is shown in each panel, with asterisks indicating statistical significance ($p < 0.05$ *, $p < 0.01$ **, $p < 0.001$***).

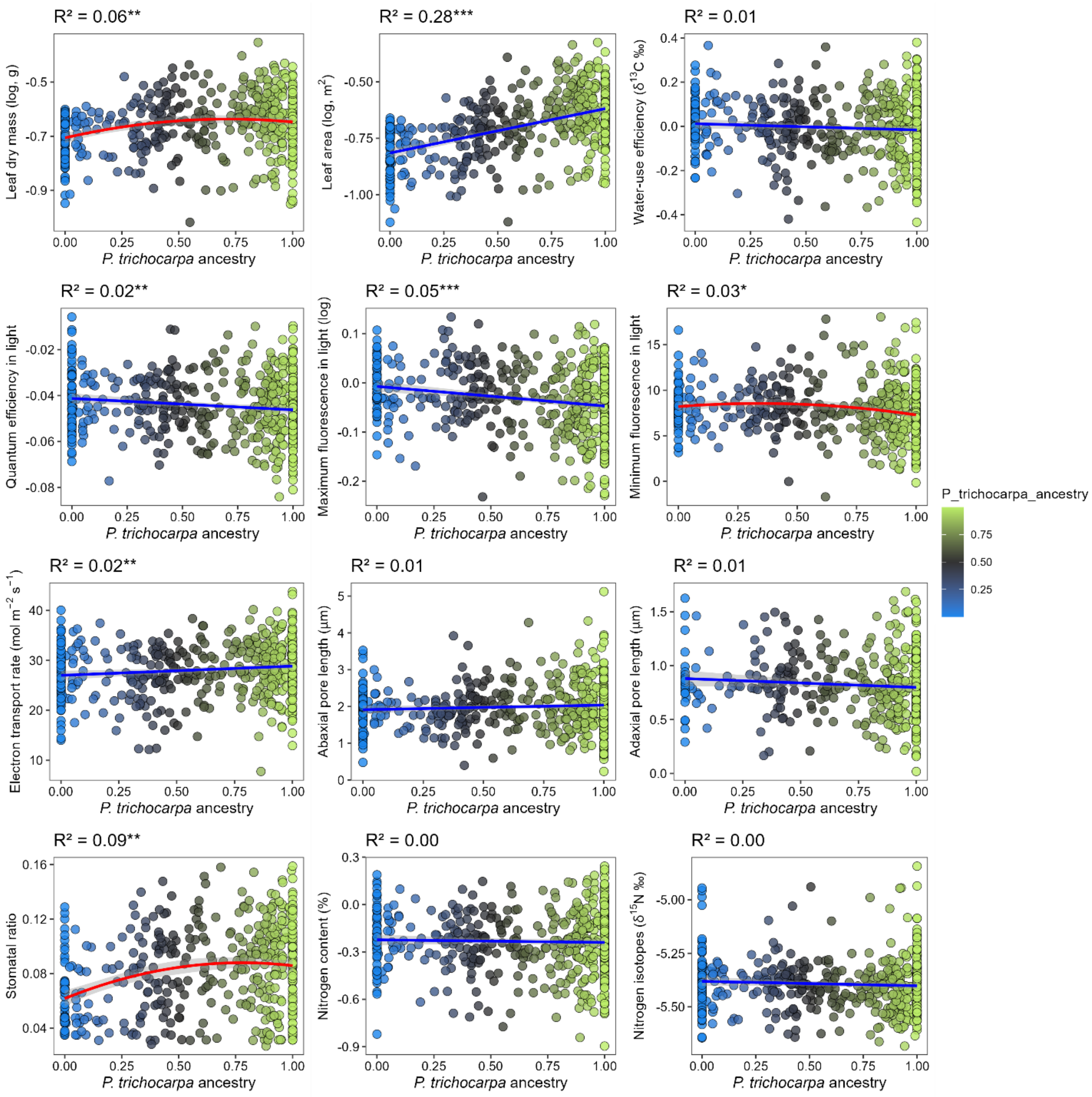


**Figure S4.** Relationship between *P. trichocarpa* ancestry (x-axis) and trait plasticity, quantified as genotype-specific reaction norm slopes (y-axis) across genotypes. Plasticity is interpreted based on the absolute magnitude of slopes: when slopes are positive, larger values indicate greater plasticity, whereas when slopes are negative, more negative values (i.e., larger absolute values) indicate greater plasticity. Points are colored by genomic ancestry, with values near 0 indicating greater *P. balsamifera* ancestry (blue), values near 1 indicating greater *P. trichocarpa* ancestry (green), and intermediate values representing admixed genotypes (gray). Solid lines represent the best-supported model for each trait, selected using AIC comparison between linear and quadratic models (ΔAIC > 2). Linear fits are shown in blue and quadratic fits in red, with shaded areas representing 95% confidence intervals. The coefficient of determination ($R^2$) for the selected model is shown in each panel, with asterisks indicating statistical significance ($p < 0.05$ *, $p < 0.01$ **, $p < 0.001$***).

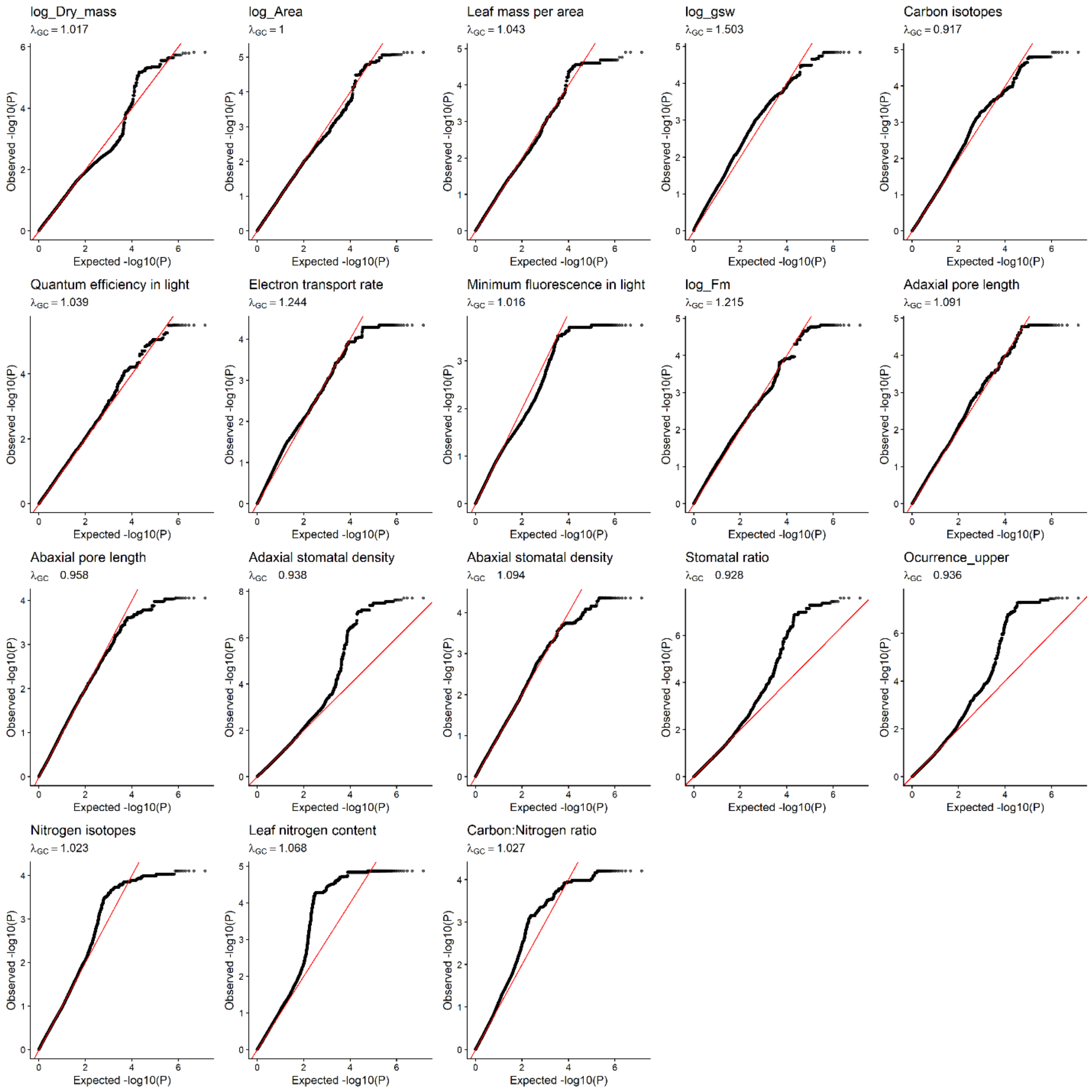


**Figure S5**. Quantile–quantile (QQ) plots of association results for trait variation in *Populus* hybrids. Each panel shows the observed versus expected –log10(P) values from GEMMA linear mixed model genome-wide association analyses. Red lines indicate the null expectation under no association. $\lambda_{GC}$ values denote the genomic control inflation factor for each trait.

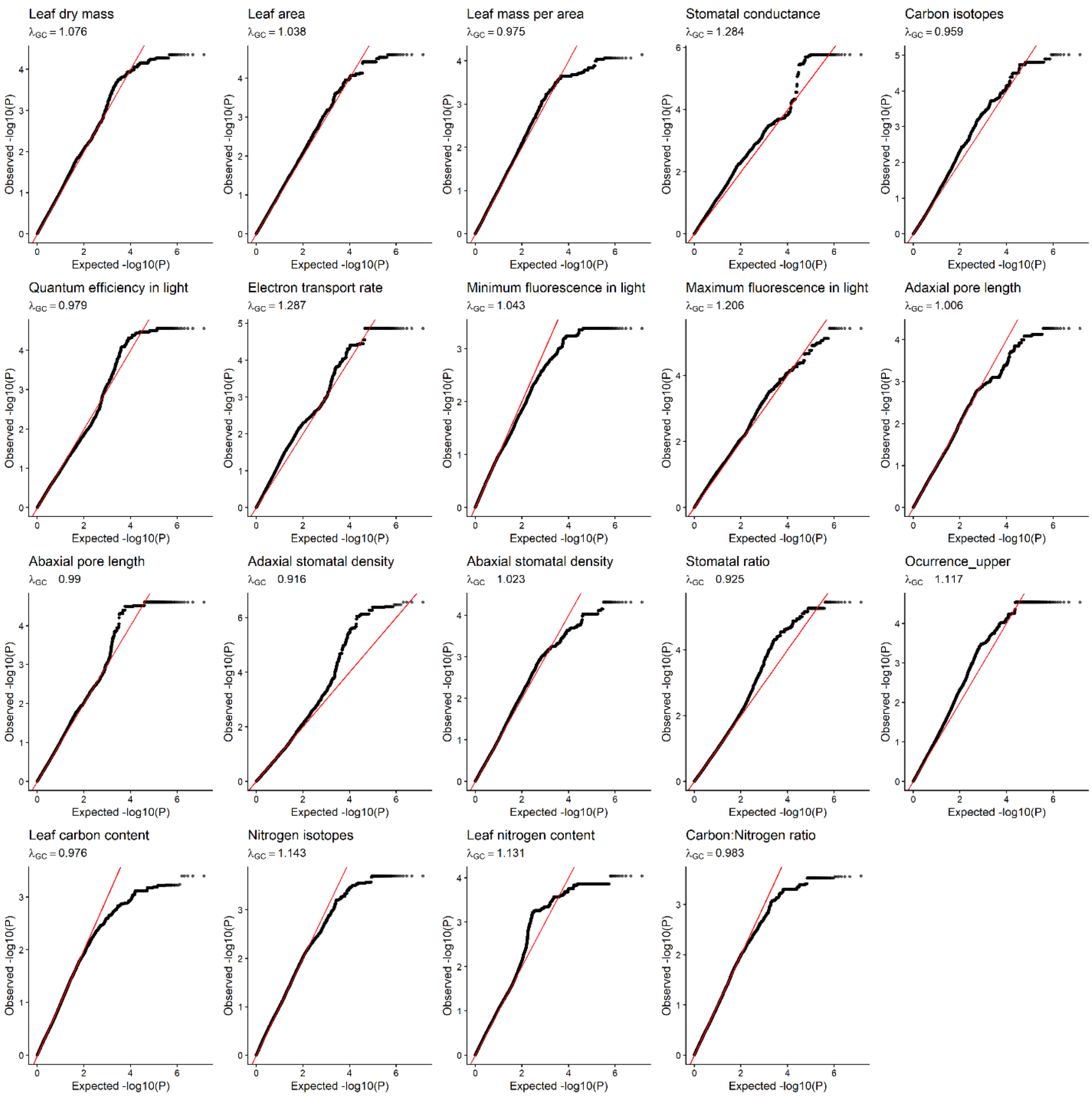


**Figure S6.** Quantile–quantile (QQ) plots of association results for slopes of reaction norms in Populus hybrids. Each panel shows the observed versus expected –log10(P) values from GEMMA linear mixed model genome-wide association analyses. Red lines indicate the null expectation under no association. $\lambda_{GC}$ values denote the genomic control inflation factor for each trait.

**Table S1**. Associations between genomic ancestry and physiological trait variation. Linear and quadratic models describing relationships between *P. trichocarpa* ancestry and physiological traits are compared, including model selection, model fit ($R^2$), model support (AIC, $\Delta$AIC), and parameter estimates (linear and quadratic slopes with associated p-values). The best-supported model was selected based on AIC, with models differing by >2 $\Delta$AIC.

| Trait variation | Best model | R2 | AIC linear model | AIC quadratic model | Delta AIC | Slope linear model | *p* value linear model | Slope quadratic model | *p* value quadratic model |
|---|---|---|---|---|---|---|---|---|---|
| ***Leaf Morphology*** | | | | | | | | | |
| Leaf dry mass (log) | linear | 0.31 | 40.96 | 42.96 | -2.00 | 0.45 | < 0.001 | -0.01 | 0.98 |
| Leaf area (log) | linear | 0.37 | 137.70 | 139.70 | -2.00 | 0.56 | < 0.001 | 0.01 | 0.98 |
| Leaf mass area | linear | 0.14 | 2815.18 | 2815.59 | -0.41 | -4.32 | < 0.001 | -5.12 | 0.21 |
| ***Gas exchange*** | | | | | | | | | |
| Stomatal conductance (log) | quadratic | 0.24 | -351.15 | -368.18 | 17.03 | -0.23 | < 0.001 | -0.73 | < 0.001 |
| Water-use efficiency | quadratic | 0.07 | 646.93 | 643.59 | 3.34 | 0.32 | < 0.001 | 1.07 | < 0.05 |
| ***Photochemistry*** | | | | | | | | | |
| Quantum efficiency in light | linear | 0.01 | -6249.28 | -6247.37 | -1.91 | 0.00 | < 0.01 | 0.00 | 0.77 |
| Maximum fluorescence in light (log) | linear | 0.04 | -3054.68 | -3053.65 | -1.03 | -0.01 | < 0.001 | -0.01 | 0.43 |
| Minimum fluorescence in light | linear | 0.02 | 1926.21 | 1925.58 | 0.64 | -0.64 | < 0.01 | -2.69 | 0.11 |
| Electron transport rate | linear | 0.01 | 2707.58 | 2706.21 | 1.37 | -0.96 | < 0.05 | -6.64 | 0.07 |
| ***Stomatal patterning*** | | | | | | | | | |
| Abaxial pore length | quadratic | 0.05 | 1607.30 | 1597.47 | 9.83 | -0.49 | < 0.001 | -4.16 | < 0.001 |
| Adaxial pore length | quadratic | 0.04 | 1118.06 | 1113.73 | 4.33 | -0.50 | < 0.01 | -2.59 | < 0.05 |
| Abaxial density | quadratic | 0.05 | 2095.38 | 2078.50 | 16.89 | 0.61 | < 0.01 | 8.45 | < 0.001 |
| Adaxial density | quadratic | 0.12 | 1720.65 | 1715.24 | 5.41 | 1.22 | < 0.001 | -3.67 | < 0.01 |
| Stomatal Ratio | quadratic | 0.12 | -1475.16 | -1488.42 | 13.26 | 0.05 | < 0.001 | -0.21 | < 0.001 |

| Trait variation | Best model | R2 | AIC linear model | AIC quadratic model | Delta AIC | Slope linear model | *p* value linear model | Slope quadratic model | *p* value quadratic model |
|---|---|---|---|---|---|---|---|---|---|
| Adaxial stomatal occurrence | quadratic | 0.13 | 1728.85 | 1713.47 | 15.38 | 1.17 | < 0.001 | -5.64 | < 0.001 |
| ***Nutrient-Use Traits*** | | | | | | | | | |
| Nitrogen content | quadratic | 0.03 | -623.47 | -636.20 | 12.73 | -0.01 | 0.72 | -0.49 | < 0.001 |
| Nitrogen isotopes | quadratic | 0.02 | -1340.20 | -1345.76 | 5.56 | -0.01 | 0.45 | -0.17 | < 0.01 |
| Carbon Nitrogen ratio | quadratic | 0.04 | 1429.93 | 1412.56 | 17.37 | 0.19 | 0.13 | 4.44 | < 0.001 |

**Table S2**. Associations between genomic ancestry and trait plasticity measured as genotype-specific reaction norm slopes. Linear and quadratic models describing relationships between *P. trichocarpa* ancestry and trait plasticity are compared, including model selection, model fit ($R^2$), model support (AIC, ΔAIC), and parameter estimates (linear and quadratic slopes with associated p-values). The best-supported model was selected based on AIC, with models differing by >2 ΔAIC.

| **Trait plasticity** | **Best model** | **R2** | **AIC linear model** | **AIC quadratic model** | **Delta AIC** | **Slope linear model** | ***p* value linear model** | **Slope quadratic model** | ***p* value quadratic model** |
|---|---|---|---|---|---|---|---|---|---|
| ***Leaf Morphology*** | | | | | | | | | |
| Leaf dry mass (log) | quadratic | 0.06 | -888.86 | -895.27 | 6.41 | 0.05 | < 0.001 | -0.28 | < 0.01 |
| Leaf area (log) | linear | 0.28 | -714.35 | -714.28 | -0.07 | 0.20 | < 0.001 | -0.16 | 0.17 |
| Leaf mass area | quadratic | 0.38 | 3696.31 | 3681.84 | 14.47 | -19.66 | < 0.001 | -39.59 | < 0.001 |
| ***Gas exchange*** | | | | | | | | | |
| Stomatal conductance (log) | quadratic | 0.24 | -1895.69 | -1908.89 | 13.20 | 0.05 | < 0.001 | 0.14 | < 0.001 |
| Water-use efficiency | linear | 0.01 | -582.31 | -580.66 | -1.65 | -0.03 | 0.08 | 0.08 | 0.56 |
| ***Photochemistry*** | | | | | | | | | |
| Quantum efficiency in light | linear | 0.02 | -2894.14 | -2892.26 | -1.88 | -0.01 | < 0.01 | -0.01 | 0.73 |
| Maximum fluorescence in light (log) | linear | 0.05 | -1308.96 | -1308.66 | -0.30 | -0.04 | < 0.001 | -0.08 | 0.19 |
| Minimum fluorescence in light | quadratic | 0.03 | 2470.67 | 2467.96 | 2.71 | -1.01 | < 0.01 | -6.19 | < 0.05 |
| Electron transport rate | linear | 0.02 | 3109.66 | 3108.08 | 1.58 | 1.85 | < 0.01 | 10.24 | 0.06 |
| ***Stomatal patterning*** | | | | | | | | | |
| Abaxial pore length | linear | 0.01 | 968.54 | 970.44 | -1.90 | 0.13 | 0.10 | -0.21 | 0.75 |
| Adaxial pore length | linear | 0.01 | 163.88 | 165.11 | -1.23 | -0.08 | 0.09 | -0.26 | 0.38 |

| Trait plasticity | Best model | R2 | AIC linear model | AIC quadratic model | Delta AIC | Slope linear model | *p* value linear model | Slope quadratic model | *p* value quadratic model |
|---|---|---|---|---|---|---|---|---|---|
| Abaxial density | quadratic | 0.03 | 1277.41 | 1267.39 | 10.03 | -0.18 | 0.08 | -2.98 | < 0.001 |
| Adaxial density | quadratic | 0.12 | 918.14 | 913.25 | 4.89 | 0.55 | < 0.001 | -1.58 | < 0.01 |
| Stomatal Ratio | quadratic | 0.09 | -2083.58 | -2091.13 | 7.55 | 0.02 | < 0.001 | -0.09 | < 0.01 |
| Occurrence upper | quadratic | 0.13 | -341.73 | -357.07 | 15.34 | 0.15 | < 0.001 | -0.71 | < 0.001 |
| ***Nutrient-Use Traits*** | | | | | | | | | |
| Nitrogen content | linear | 0.00 | -205.36 | -206.71 | 1.35 | -0.02 | 0.49 | 0.36 | 0.07 |
| Nitrogen isotopes | linear | 0.00 | -667.85 | -669.12 | 1.28 | -0.02 | 0.16 | 0.22 | 0.07 |
| Carbon Nitrogen ratio | linear | 0.00 | 1394.63 | 1396.03 | -1.40 | 0.06 | 0.64 | -0.76 | 0.44 |

**Table S3**. List of candidate genes underlying physiological trait variation and plasticity identified through admixture mapping analysis.

| Trait | Poplar gene | Arabidopsis | Method | Description |
|---|---|---|---|---|
| Stomatal Ratio | Potri.005G103700 | AT4G35690 | Reaction norms | Arabidopsis protein of unknown function |
| Stomatal Ratio | Potri.005G104700 | AT3G51390 | Reaction norms | Belongs to the DHHC palmitoyltransferase family |
| Stomatal Ratio | Potri.009G012200 | AT5G60490 | Trait mean | Fasciclin-like arabinogalactan protein |
| Stomatal Ratio | Potri.009G012300 | AT1G07980 | Trait mean | DNA polymerase epsilon subunit |
| Stomatal Ratio | Potri.009G014600 | AT2G28370 | Reaction norms | Belongs to the Casparian strip membrane proteins (CASP) family |
| Stomatal conductance | Potri.010G224100 | AT2G37040 | Reaction norms | Phenylalanine ammonialyase |
| Stomatal conductance | Potri.010G224200 | AT2G37040 | Reaction norms | Phenylalanine ammonialyase |
| Stomatal conductance | Potri.010G224300 | AT5G03300 | Reaction norms | adenosine kinase |
| Adaxial stomatal density | Potri.012G038300 | AT1G18330 | Trait mean | transcription, DNA-templated (Early-phytochrome-responsive1) |
| Stomatal Ratio | | | Trait mean, Reaction norms | |
| Adaxial stomatal occurrence | Potri.015G022000 | AT5G53200 | Trait mean, Reaction norms | transcription factor TWIST |
| Adaxial stomatal density | | | Trait mean, Reaction norms | |
| Stomatal Ratio | | | Trait mean | |
| Adaxial stomatal occurrence | Potri.015G022100 | AT5G24120 | Trait mean, Reaction norms | RNA polymerase sigma factor sigE, chloroplastic |
| Adaxial stomatal density | | | Trait mean | |
| Stomatal Ratio | | | Trait mean, Reaction norms | |
| Adaxial stomatal occurrence | Potri.015G022300 | AT5G53210 | Trait mean, Reaction norms | Transcription factor TRY |
| Adaxial stomatal density | | | Trait mean | |

| Trait | Poplar gene | Arabidopsis | Method | Description |
|---|---|---|---|---|
| Stomatal Ratio | | | Trait mean, Reaction norms | |
| Adaxial stomatal occurrence | | | Trait mean, Reaction norms | |
| Stomatal Ratio | Potri.015G022400 | AT5G41480 | Trait mean | Dihydrofolate synthetase (DFA) |
| SDR | | | Trait mean, Reaction norms | |
| Adaxial stomatal occurrence | Potri.015G022500 | AT5G53220 | Trait mean, Reaction norms | hypothetical protein;(source:Araport11) |
| Stomatal Ratio | | | Trait mean | |
| Adaxial stomatal occurrence | Potri.015G022600 | AT5G53250 | Trait mean, Reaction norms | Arabinogalactan peptide (AGP22) |
| Adaxial stomatal density | | | Trait mean | |
| Adaxial stomatal occurrence | Potri.015G022700 | AT4G27940 | Trait mean, Reaction norms | Belongs to the mitochondrial carrier (TC 2.A.29) family (MTM1) |
| Adaxial stomatal density | | | Trait mean | |
| Adaxial stomatal occurrence | Potri.015G022800 | AT5G43360 | Trait mean, Reaction norms | Inorganic phosphate transporter (PHT3,1) |
| Adaxial stomatal density | | | Trait mean | |
| Adaxial stomatal occurrence | Potri.015G022900 | AT5G53280 | Trait mean, Reaction norms | Plastid division protein (PDV1) |
| Adaxial stomatal density | | | Trait mean | |
| Adaxial stomatal occurrence | Potri.015G023000 | AT1G14820 | Trait mean, Reaction norms | CRAL/TRIO domain |
| Adaxial stomatal density | | | Trait mean | |
| Leaf area | Potri.017G115700 | AT1G67050.2 | Trait mean | - |
| Leaf dry mass | | | Trait mean | |
| Leaf dry mass | Potri.017G115801 | AT1G67050.3 | Trait mean | - |
| Leaf dry mass | Potri.017G115900 | AT3G47580 | Trait mean | Belongs to the protein kinase superfamily. Ser Thr protein kinase family |
